# Mast cell desensitization induces a distinct IgE-dependent transcriptional program associated with immune regulation

**DOI:** 10.64898/2026.02.28.708745

**Authors:** Celia López-Sanz, Emilio Nuñez-Borque, Alessandra Ruiz-Sánchez, Javier Sevilla-Montero, Antonio J. García-Cívico, Sofía Álvarez-Garrote, Marcos Zamora-Dorta, Elisa Sánchez-Martínez, Lucía Moreno-Serna, Maricruz Mamani-Huanca, Alma Villaseñor, Domingo Barber, Eduardo Balsa, Cecilia Muñoz-Calleja, Pedro Ojeda, Carlos Blanco, Pablo Rodríguez del Río, Francisco Sánchez-Madrid, Rodrigo Jiménez-Saiz

## Abstract

Allergen-driven IgE–mast cell (MC) activation is a central feature of allergic diseases, whose prevalence continues to increase worldwide. Allergen immunotherapy (AIT) is currently the only disease-modifying treatment and induces a state of MC hyporesponsiveness termed desensitization; however, its underlying molecular mechanisms remain incompletely understood and whether this state reflects passive signal attenuation or active cellular reprogramming remains unresolved. Here, we define the molecular landscape of MC desensitization using a human polyclonal platform that captures the physiological diversity of allergen-specific IgE. Desensitization reduced degranulation in an allergen-specific manner and induced progressive internalization of allergen-specific IgE. Although early steps were associated with LAT phosphorylation, subsequent allergen challenge failed to propagate activation to distal IgE/FcεRI effectors, revealing selective signaling uncoupling. Transcriptomic profiling uncovered a distinct transcriptional program comprising 168 upregulated genes enriched in immunoregulatory pathways and largely non-overlapping with classical activation signatures. This reprogramming occurred despite minimal alterations in mitochondrial respiration and selective impairment of allergen-induced glycolysis. Functionally, desensitized MCs enhanced allergen-driven proliferation of memory CD4^⁺^ T cells. Together, these findings demonstrate that MC desensitization is not merely passive hyporesponsiveness but involves time-dependent allergen-specific IgE internalization, selective signal propagation, and a unique immunoregulatory transcriptional imprint that may contribute to tolerance during AIT.

## INTRODUCTION

In the last decades, allergic diseases have reached near-pandemic proportions, affecting more than 20% of the global population and imposing a substantial burden on healthcare systems and the economy^1, 2^. Although heterogeneous in clinical presentation, allergic diseases are characterized by exacerbated type 2 immune responses to otherwise harmless antigens, driven by immunoglobulin (Ig)E and effector cells, such as basophils and mast cells (MCs)^3, 4^. In sensitized individuals, allergen-mediated crosslinking of FcLRI-bound IgEs on MCs activates Src family kinases (Lyn and Fyn), leading to the phosphorylation of immunoreceptor tyrosine-based activation motifs, the recruitment of Syk and LAT, and the subsequent engagement of downstream events, including PLCγ, PI3K-Akt and MAPK-ERK signaling^5^. Ultimately, these pathways culminate in rapid degranulation and release of preformed inflammatory mediators that cause clinical manifestations related to allergic diseases, ranging from mild to life-threatening reactions, including anaphylaxis^6, 7, 8^.

Allergy management relies largely on allergen avoidance and pharmacological symptom control. However, accidental allergen exposure remains common, particularly in food allergy^9, 10, 11^, and environmental allergens are often unavoidable. Conventional drugs do not provide sustained disease modification and allergen immunotherapy (AIT) is currently the only specific intervention with disease-modifying potential^12, 13^. AIT has proven efficacy for certain allergies, such as Hymenoptera venom allergy^14^ or allergic rhinitis for major allergens, but weaker evidence for others^13^. For food allergies, oral AIT has emerged in the last two decades and constitutes now an accepted treatment option^15^. However, AIT still faces several limitations. For example, AIT remains experimental for various indications, which restricts its broader implementation^16^. Additionally, adverse reactions, including anaphylaxis, can occur during the treatment, compromising patients’ safety and adherence^17, 18, 19, 20^.

AIT begins with a build-up phase, during which increasing doses of the allergen are administered until the target dose is reached. Paradoxically, during this build-up phase, although patients exhibit increased IgE levels, MCs become unresponsive to the allergen (desensitized)^21^. Subsequently, during the maintenance phase, the target allergen dose is administered at regular intervals. This phase is characterized by a gradual decline in allergen-specific IgE levels while IgG4 and IgA levels increase due to changes in different sub-populations of B– and CD4^+^ T-cells^18, 22, 23, 24, 25^. Effector cell desensitization is a critical determinant of AIT safety, as it limits treatment-associated adverse reactions. Moreover, effector-cell desensitization has been linked to sustained unresponsiveness in food-allergic patients^26^ and implicated in the establishment of oral tolerance in food-allergy models of AIT^27^. Notwithstanding, the molecular mechanisms underlying desensitization remain poorly defined and, in some aspects, controversial^21^.

Conflicting reports have proposed divergent mechanisms. Oka *et al*. described the allergen-specific IgE internalization as a central feature of MC desensitization^28^. In sharp contrast, other studies suggested that IgE/FcεRI internalization is limited during desensitization and instead proposed impaired calcium flux and actin remodeling as key determinants^29^. More recently, Adnan *et al.* reported the absence of IgE/FcεRI internalization during desensitization and proposed that this process is partially regulated by the phosphatase SHIP-1^30^. These contrasting results evidence mechanistic discrepancies, particularly regarding IgE turnover and proximal signaling. Importantly, most studies have relied on monoclonal IgE models, which may not fully recapitulate the complexity of the human polyclonal IgE repertoire^31, 32^. Furthermore, beyond IgE internalization, it is unknown whether desensitization induces broader cellular changes, such as metabolic or transcriptomic shifts, that could influence long-term immunomodulation during AIT.

Here, we investigate the molecular basis of MC desensitization using a polyclonal desensitization platform^33^. We dissect the dynamics of IgE/FcLRI internalization and downstream signaling and evaluate the metabolic and transcriptomic consequences of desensitization. We show that desensitization induces an allergen-specific, time-dependent IgE internalization that limits allergen responsiveness while preserving selective IgE/FcLRI signaling. By RNA-sequencing (RNA-seq) and RT-qPCR analyses, we reveal that desensitized MCs acquire a distinct transcriptional program enriched in immunoregulatory pathways and demonstrate that these changes influence CD4^+^ T-cell responses in an allergic context.

## RESULTS

### Time dependent internalization of allergen-specific IgE during desensitization limits MC degranulation

To ascertain the intrinsic mechanisms involved in MC desensitization, we used a previously reported polyclonal protocol^33^. Since this protocol was optimized for peanut (PN) allergens, we first validated whether the same experimental conditions (*i.e*., desensitization steps, dose and time intervals) also worked using sera from patients allergic to other foods (egg and milk) and environmental allergens (pollen and cat dander). Therefore, we compared human LAD2 MCs sensitized (ALL) with sera from allergic donors (**Table S1**) and desensitized (DS) and evaluated MC degranulation by measuring CD63 and CD107a expression (**Fig. S1a**) along with β-hexosaminidase. Following desensitization, we observed a significant reduction of degranulation in DS compared to ALL cells upon allergen stimulation, indicating that the protocol was neither serum nor allergen exclusive (**Fig. 1a**). Meanwhile, provocation with the positive control (anti-IgE) recovered MC degranulation, with no differences between ALL and DS cells, indicating intact degranulation machinery. To validate these findings, we repeated the protocol in primary human MCs (hMCs) derived from peripheral blood CD34^+^ progenitors and sensitized with the same allergic sera. Although we observed an increase in degranulation during desensitization in hMCs, the cells remained unresponsive to allergen challenge, with no differences in DS cells between challenged and unchallenged conditions (**Fig. 1b**). These data suggest that the mechanisms operating in this polyclonal desensitization protocol extend to different types of human MCs and IgE repertoires.

**Figure 1.**
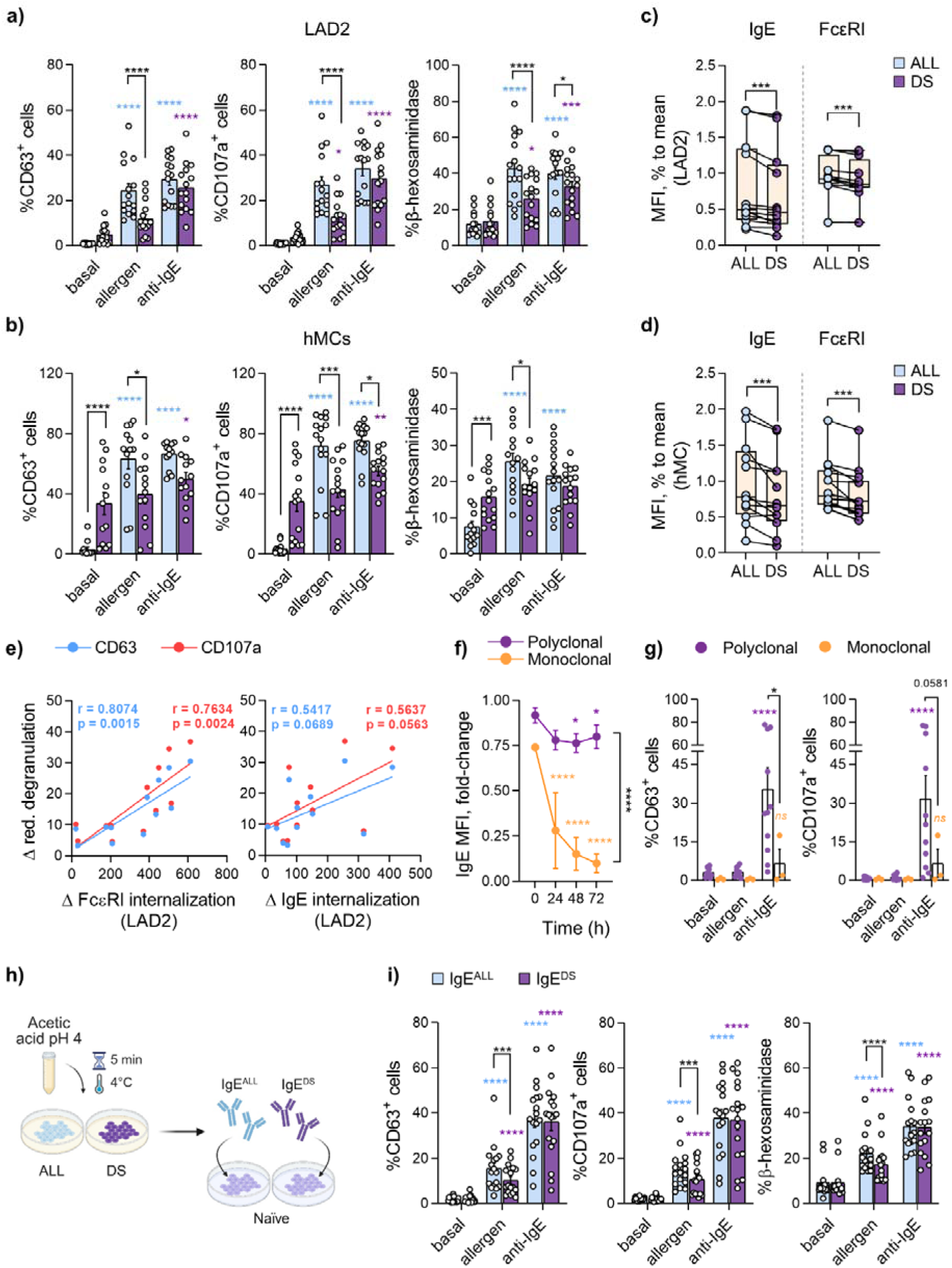
Time-dependent and allergen-specific IgE internalization during desensitization. Mast cells (MCs) were sensitized with sera from patients allergic to peanut (n=3), egg white (n=3), milk (n=3), pollen (n=2-3) and cat (n=2-3). **(a, b)** CD63 and CD107a expression and β-hexosaminidase activity after allergen or anti-human (h)IgE provocation (20 µg/mL) in allergic (ALL) and desensitized (DS) LAD2 cells **(a)** or primary hMCs **(b)**. **(c, d)** IgE and FcεRI mean fluorescence intensity (MFI) quantification in ALL and DS LAD2 **(c)** and hMCs **(d)** once the protocol was completed. MFI values were normalized to the mean of ALL values. **(e)** Pearson correlation between IgE and Fc□RI internalization (Δ internalization: MFI ALL-MFI DS) and CD63 (blue) and CD107a (red) reduction (Δ reduction of degranulation; % degranulation ALL – % degranulation DS) in LAD2 cells. Pearson r and p-values for each degranulation marker are shown in matching colors; (n=13). **(f)** Kinetic evaluation (0, 24, 48 and 72 h) of IgE MFI in polyclonal (purple; n=8) or monoclonal (orange; n=3) DS LAD2 or bone-marrow derived MCs (BMMCs), respectively. ALL BMMCs were sensitized with ovalbumin (OVA)-specific IgE (clone EC1). **(g)** Degranulation (CD63 and CD107a) was evaluated 72 h post-desensitization following allergen (20µg/mL) or OVA (10 µg/mL) and anti-hIgE (20 µg/mL) or anti-murine IgE (10 µg/mL) provocations. **(h)** Experimental design. IgEs from ALL or DS LAD2 cells were stripped using mild acetic acid wash (pH 4). Stripped ALL (IgE^ALL^) and DS (IgE^DS^) IgEs were used to sensitize naïve LAD2 cells. Image created with Biorender.com. **(i)** CD63 and CD107a expression and β-hexosaminidase activity upon allergen and anti-hIgE (20 µg/mL) provocation in IgE^ALL^ or IgE^DS^ sensitized LAD2 cells; (n=17). Data are represented as mean ± SEM and box plots with median and interquartile range. Comparisons between ALL *vs.* DS (c, d) were made with paired Wilcoxon test. Multiple comparisons between groups (a, b, f, g, i) were made using two-way ANOVA, paired by serum, followed by Bonferroni correction test. Data that did not follow normal distribution (b, h) was logarithmically transformed before two-way ANOVA was performed. Colored asterisks indicate significant differences relative to matched basal controls. *p<0.05, **p<0.01, ***p<0.001, ****p<0.0001.

Given that IgE internalization has been linked to MC desensitization in monoclonal protocols^28, 29^, we evaluated the internalization of IgE/FcεRI complexes under the polyclonal protocol by flow cytometry. A significant reduction of IgE and FcεRIα levels was detected in DS cells in both LAD2 and hMCs (**Fig. 1c-d, Fig. S2a**). Accordingly, we found a positive correlation between the degree of FcεRI internalization and the reduction in degranulation in LAD2 cells (**Fig. 1e, left**), with a slightly weaker correlation for IgE (**Fig. 1e, right**). These findings prompted us to better define the IgE internalization dynamics; thus, we analyzed IgE levels over 72 h following polyclonal desensitization. IgE levels declined immediately after desensitization, reaching a statistically significant nadir at 48 h and remaining low for up to 72 h (**Fig. 1f**). Notably, although IgE was still detectable at the endpoint, it was unable to induce allergen-specific degranulation on MCs while retaining responsiveness to anti-IgE stimulation (**Fig. 1g, purple**). These results suggested that only the allergen-specific IgE was being internalized during desensitization, while the non-specific IgE remained on the surface of MCs. To confirm these results, we studied IgE dynamics using the Oka *et al*.^28^ ovalbumin (OVA)-monoclonal model in bone-marrow derived MCs (BMMCs). We observed that IgE internalization was faster than under polyclonal conditions. Importantly, BMMCs failed to respond to either allergen or anti-IgE challenge 72 h after desensitization (**Fig. 1g, orange**), indicating that desensitization was presumably driven by specific IgE internalization.

To test whether surface-bound IgE persisting after desensitization was indeed non-specific, we stripped IgE from ALL and DS LAD2 cells and used these recovered IgEs to sensitize naïve MCs (**Fig. 1g, Fig. S2b**). Upon challenge, IgE from DS cells (IgE^DS^) induced lower MC degranulation compared to the IgE taken from ALL controls (IgE^ALL^) (**Fig. 1i**). Conversely, re-sensitization of stripped ALL and DS MCs with fresh allergic sera restored full degranulation upon stimulation (**Fig. S2c-d**), confirming that desensitization does not compromise the degranulation capacity of MCs. Altogether, these results indicate that desensitization induces IgE internalization in an allergen-specific manner, which limits MC activation upon subsequent allergen encounter.

### Desensitization induces selective signal transduction in MCs and prevents the onset of degranulation

The allergen-specific internalization of IgE during desensitization is consistent with ongoing IgE-mediated signaling. Previous studies using monoclonal desensitization models focused on individual components of the IgE/FcLRI pathway^30, 34^, but did not evaluate their coordinated regulation. Therefore, we systematically examined proximal (Lyn, LAT, SHIP-1) and distal (PLCγ, ERK, Akt) IgE/FcLRI signaling intermediates (**Fig. S3a**) across desensitization steps and compared them with ALL MCs challenged with matched allergen concentrations (**Fig. 2a, b**). Among all the upstream proteins, an earlier partial phosphorylation of SHIP-1 was observed during desensitization steps compared to single-challenge controls (**Fig. 2c**), supporting the role of this phosphatase in desensitization^30^. However, no significant differences were found in the phosphorylation of the activating (Y416) and inhibitory (Y507) residues of Lyn (**Fig. 2d**), nor in their ratio (**Fig. S3b**).

**Figure 2.**
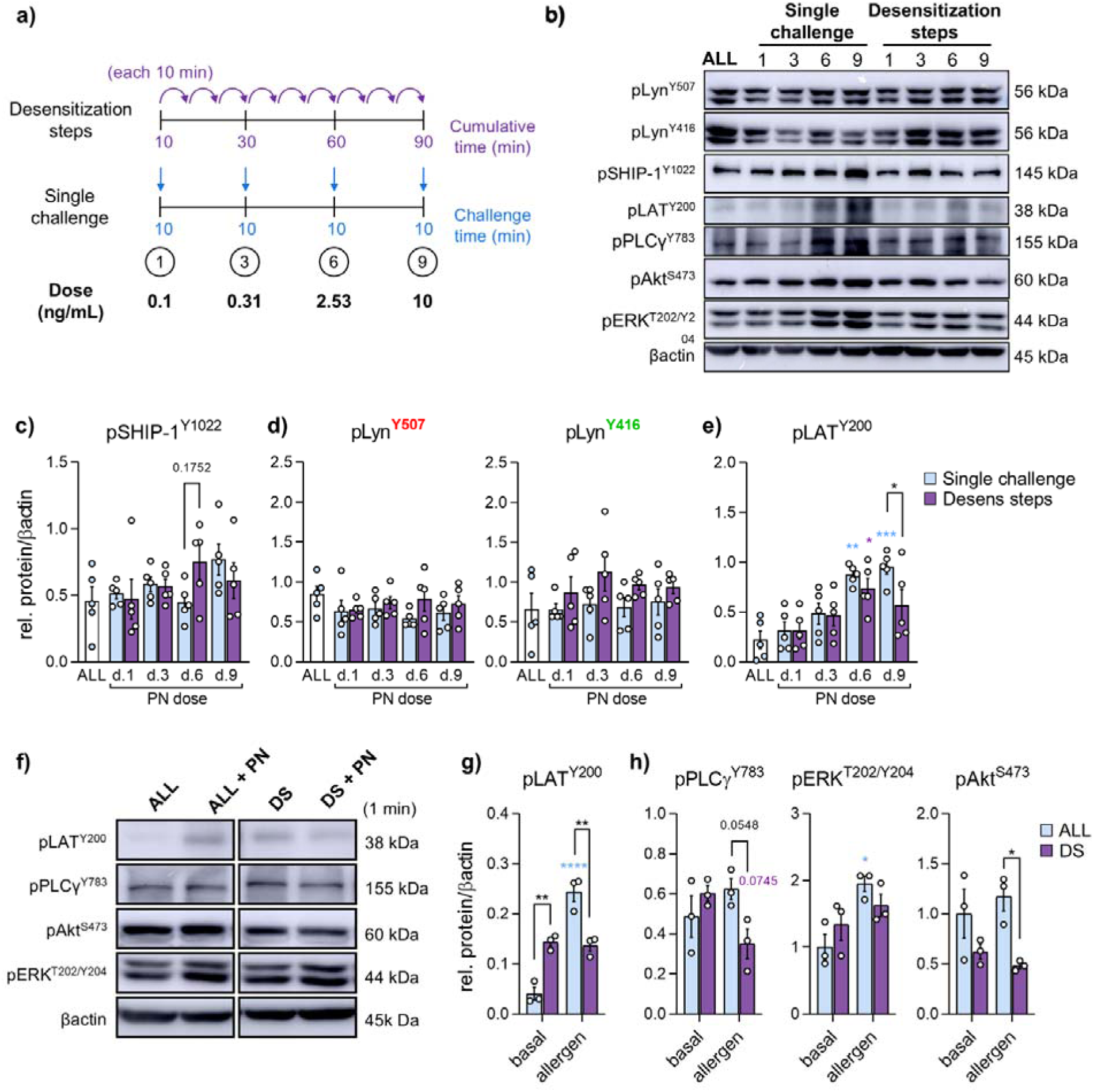
IgE/FcεRI signaling pathways triggered during mast cell desensitization. **(a)** Experimental design. LAD2 cells were sensitized with peanut (PN)-allergic sera and either desensitized or stimulated with single-dose PN challenges matching the cumulative allergen dose reached at each desensitization step (0.1, 0.31, 2.53, and 10□ng/mL). **(b)** Representative Western blot showing phosphorylation of proximal (Lyn, LAT, SHIP-1) and distal (PLCγ, Akt, ERK) components of the IgE/FcεRI signaling cascade. β-actin was used as a loading control. **(c–e)** Quantification of SHIP-1 (Y1022) **(c)**, Lyn (Y507) and Lyn (Y416) **(d)**, and LAT (Y200) **(e)** phosphorylation relative to β-actin (n□=□5). **(f)** Representative Western blot showing phosphorylation of LAT, PLCγ, ERK, and Akt in PN-ALL and DS LAD2 cells following 1□min stimulation with PN (20□µg/mL). **(g, h)** Quantification of LAT (Y200) **(g)** PLCγ (Y783), ERK (T202/Y204) and Akt (S473) phosphorylation **(h)** relative to β-actin (n□=□3). Data are presented as mean ± SEM. Statistical comparisons were performed using two-way ANOVA followed by Bonferroni multiple comparison test. Colored asterisks indicate significant differences relative to matched basal controls. *p□<□0.05.

Since LAT phosphorylation at Y226 residue has been previously reported in ALL and DS cells^34^, we next evaluated phosphorylation at the Y200 residue within the C-terminal regulatory domain. Interestingly, escalating allergen doses induced a progressive increase in LAT phosphorylation, whereas desensitization resulted in a significant attenuation at step 9 compared to the corresponding ALL condition (**Fig. 2e**). These changes in LAT phosphorylation paralleled those in MC degranulation (**Fig. S3c**) and suggested partial uncoupling between LAT activation and downstream signaling events. Accordingly, the phosphorylation pattern of PLCγ mirrored the LAT profile, exhibiting a modest reduction in DS cells at step 9 compared to its ALL homologue (**Fig. S3d**). In contrast, ERK or Akt phosphorylation remained unchanged (**Fig. S3e**).

To further assess the attenuation of IgE/FcLRI signaling following desensitization, we quantified phosphorylation of proximal and distal intermediates in ALL and DS cells after 1 min stimulation with 20 µg/mL of allergen (**Fig. 2f. Fig. S3f**). As previously observed, basal LAT phosphorylation was increased in DS cells compared with ALL controls (**Fig. 2e, g**), demonstrating active signal transduction during desensitization. Notably, compared to ALL cells, LAT phosphorylation did not further increase in DS cells following allergen challenge (**Fig. 2g**). Consistent with the LAT profile, downstream signaling intermediates PLCγ, ERK and Akt displayed reduced activation upon allergen challenge in DS cells compared with ALL cells (**Fig. 2h**), confirming that canonical IgE-mediated MC degranulation pathways are not fully engaged following desensitization. In contrast, phosphorylation of Lyn and SHIP-1 remained unchanged (**Fig. S3g**). Collectively, these findings revealed that polyclonal MC desensitization elicits selective signal transduction, while limiting activation of the principal degranulation pathways.

### Desensitization impairs allergen-induced glycolytic responses in MCs

Recent studies in monoclonally-sensitized BMMCs have shown that MCs rapidly undergo a metabolic shift toward glycolysis following IgE-dependent activation^35, 36^. Moreover, pharmacological inhibition of glycolysis alleviates allergic responses in a murine model of food allergy^37^, suggesting that glycolytic metabolism influences MC function. To determine whether desensitization alters MC metabolism, we performed metabolomic profiling of supernatants of ALL and DS LAD2 upon completion of the desensitization protocol. To minimize potential artifacts arising from serum components, we sensitized LAD2 cells with myeloma-IgE and desensitized using an anti-IgE protocol^33^. A total of 91 metabolites were detected across ALL and DS samples, although no statistically significant differences were found between conditions. In agreement with these data, neither unsupervised principal component analysis (PCA; **Fig. 3a**) nor hierarchical clustering (**Fig. S4a**) demonstrated clear separation between groups, indicating that desensitization did not induce rapid changes in secreted or consumed metabolites.

**Figure 3.**
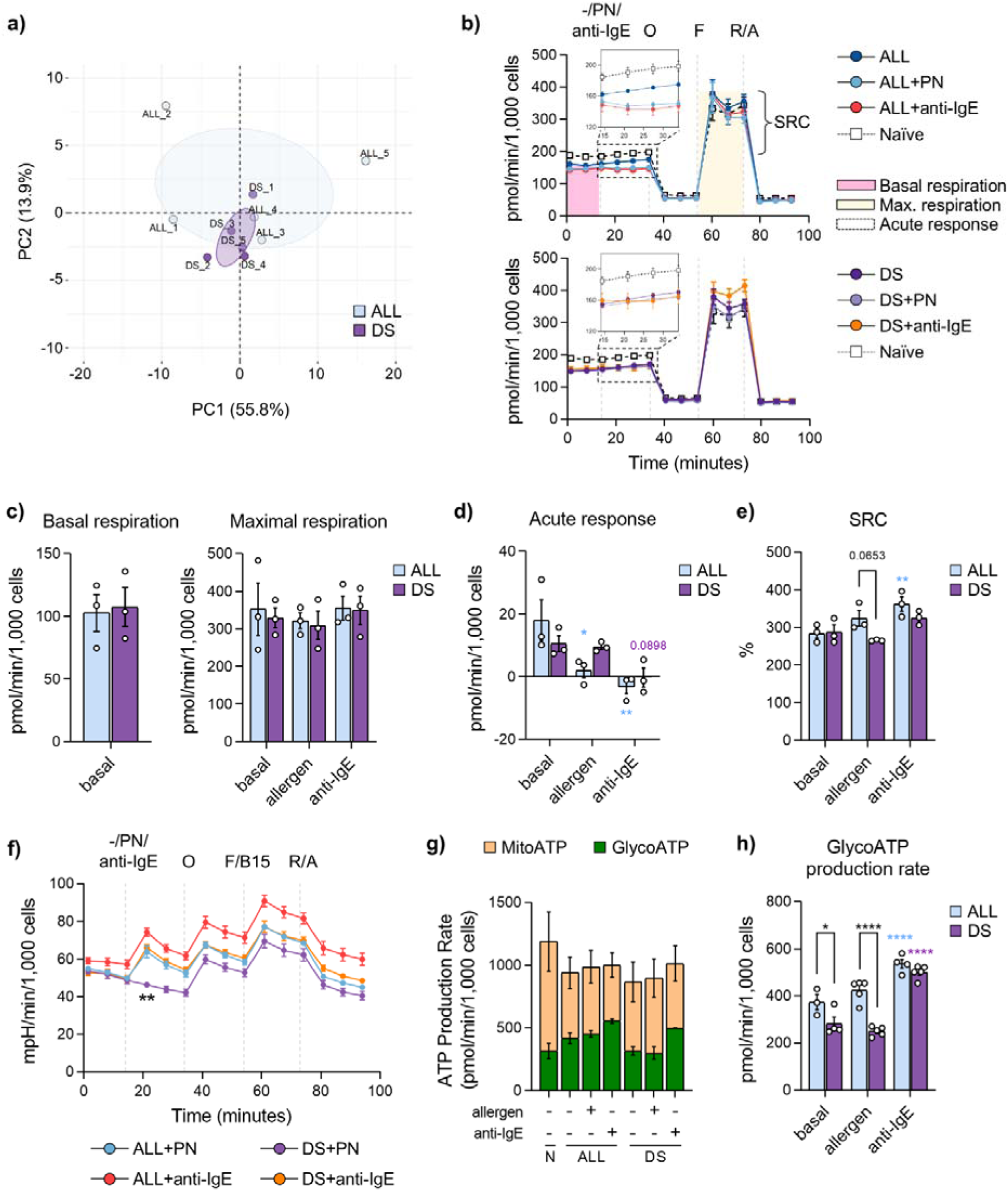
Metabolic parameter of mast cells following desensitization. **(a)** Principal component analysis (PCA) of 91 metabolites identified by mass spectrometry in supernatants from sensitized (ALL) and desensitized (DS) LAD2 cells (n□=□5). ALL cells were sensitized with human (h) myeloma IgE (100□ng/mL) and desensitized using a 7-step anti-hIgE protocol. Metabolomic profiling was performed immediately after completion of desensitization. **(b–h)** Mitochondrial respiration and glycolytic function were assessed in peanut (PN)–sensitized ALL and DS LAD2 cells using the Seahorse Mito Stress Test. **(b)** Oxygen consumption rate (OCR) traces in ALL (upper panel) and DS (lower panel) cells under basal conditions and following PN (20□µg/mL) or anti-hIgE (10□µg/mL) stimulation. Insert figure represents the acute response following PN or anti-hIgE injections. **(c–e)** Quantification of OCR-derived basal and maximal respiration **(c)**, acute OCR response to stimulation **(d)**, and spare respiratory capacity (%) **(e)** (n□=□3 independent experiments, each performed in quintuplicate). **(f)** Extracellular acidification rate (ECAR) under basal conditions and following PN (20□µg/mL) or anti-hIgE (10□µg/mL) stimulation (n□=□3 independent experiments, each in quintuplicate). **(g)** Induced ATP production rate. Quantification of mitochondrial and glycolytic ATP production rates; glycolytic ATP data are representative of two independent experiments (n□=□2 independent experiments, each in quintuplicate). **(h)** Relative contribution of glycolytic ATP production under basal and stimulated conditions. Data were normalized to cell number and are presented as mean ± SEM. Statistical comparisons were performed using two-way ANOVA followed by Bonferroni multiple comparison test (c–h). For ECAR analysis (f), comparisons were made relative to the ALL□+□PN condition across the indicated time points following drug injections (1, 20, 40, 60, and 80□min). Colored asterisks indicate significant differences relative to matched basal controls. *p□<□0.05; **p□<□0.01; ****p□<□0.0001. PC = principal component; SRC = spare respiratory capacity; O, oligomycin; F, FCCP; B15, BAM15; R/A, rotenone/antimycin A.

We next assessed the intracellular metabolic response of MCs by quantifying oxygen consumption rate (OCR) (**Fig. 3b**), a readout of mitochondrial aerobic respiration, along with OCR-derived metabolic parameters. ALL and DS cells exhibited comparable values of basal and maximal respiration (**Fig. 3c**), as well as similar proton leak, non-mitochondrial respiration and coupling efficiency (**Fig. S4b**). Remarkably, OCR decreased in ALL cells following challenge, whereas DS cells displayed no significant changes (**Fig. 3b, 3d**). In addition, the spare respiratory capacity, a marker of metabolic fitness, did not increase in DS cells following allergen challenge, but remained unchanged under anti-IgE stimulation or basal conditions (**Fig. 3e**). Overall, these results suggest an impaired allergen-specific responsiveness after desensitization.

Simultaneously, we evaluated the extracellular acidification rate (ECAR), a surrogate marker of glycolytic activity. Following allergen stimulation, ECAR increased in ALL cells but not in DS cells, whereas anti-IgE stimulation enhanced glycolysis in both conditions (**Fig. 3f**), confirming this unresponsiveness after desensitization. To further assess the functional impact of desensitization on cellular bioenergetics, we quantified the relative contribution of mitochondrial *vs*. glycolytic ATP production. No differences were observed between ALL or DS cells at the basal state (**Fig. S4c**); hence, subsequent analyses focused on stimulated states. While mitochondrial ATP production was comparable between groups (**Fig. 3g**), glycolytic ATP production was reduced in DS cells under both basal and allergen-challenged conditions but was restored following anti-IgE stimulation (**Fig. 3h, Fig. S4d**). Collectively, these data indicate that desensitization selectively blunts allergen-induced metabolic reprogramming in MCs, characterized by impaired OCR and glycolytic responses despite preserved basal mitochondrial function. This metabolic hyporesponsiveness is stimulus-specific, as anti-IgE stimulation restores glycolytic activity and ATP production in DS cells.

### Desensitization induces a distinct transcriptomic program in MCs

To more comprehensively assess the impact of desensitization on MCs, we isolated RNA from four biological replicates of naïve, PN-ALL, and DS cells and performed paired-end RNA-seq analysis 2 h after completion of the desensitization protocol (**Fig. S5a**). Unsupervised PCA of the 13,168 identified genes clustered together and separated among themselves the three experimental conditions, distinguishing mainly the DS cells from the other groups (**Fig. 4a**). Consistent with this segregation, only 17 differentially expressed genes (DEGs) were identified between ALL and naïve control cells (adjusted *p* > 0.05 and |log_2_FC| ≥ 1; **Fig. 4b, left**), whereas 187 DEGs were detected between DS and ALL cells, including 168 upregulated and 19 downregulated genes (**Fig. 4b, right**). Hierarchical clustering based on the transcriptional data from these 187 DEGs further separated DS and ALL samples into two distinct groups (**Fig. 4c**). Together, these results indicate that sensitization alone does not substantially remodel the MC transcriptome, whereas desensitization induces a marked transcriptional reprogramming.

**Figure 4.**
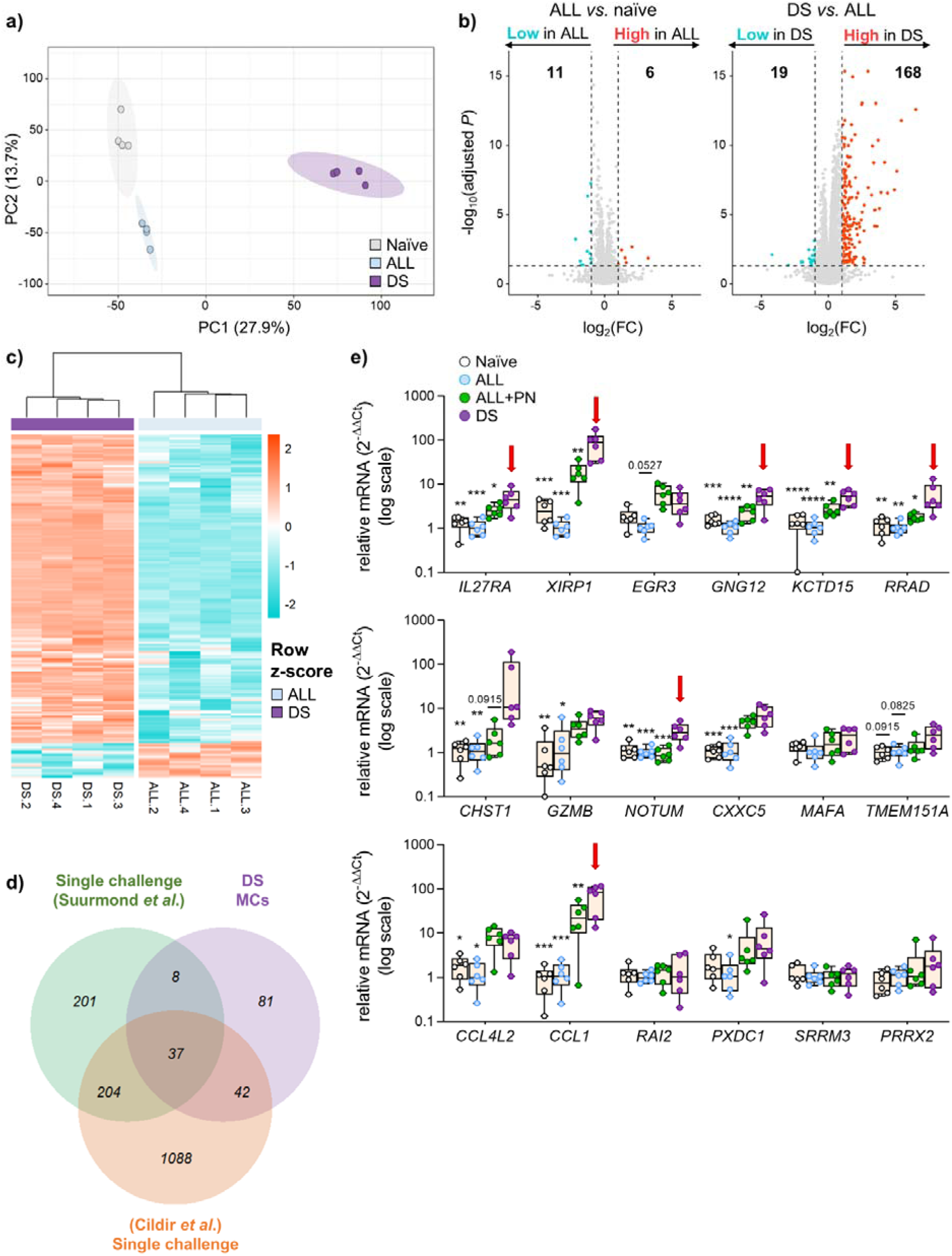
Desensitized mast cells present a distinct transcriptomic signature. RNA from four biological replicates of naïve, peanut (PN)-sensitized (ALL), and desensitized (DS) LAD2 cells was analyzed by RNA sequencing (RNA-seq) 2□h after completion of desensitization. **(a)** Principal component analysis (PCA) of 13,168 expressed genes in naïve, ALL, and DS cells. **(b)** Volcano plots comparing ALL *vs*. naïve (left) and DS *vs*. ALL (right) groups. Genes with adjusted p□<□0.05 and |log₂(FC)|□≥□1 were considered differentially expressed genes (DEGs). Blue, significantly downregulated; red, significantly upregulated; gray, not significant. **(c)** Heatmap showing hierarchical clustering based on the 187 DEGs identified between DS and ALL cells. Rows are scaled by gene expression. **(d)** Venn diagram showing overlap between upregulated DEGs in DS cells and genes reported in single-challenged human mast cells (hMCs) from published datasets: (i) Suurmond *et al*., hMCs stimulated for 6□h with anti-hIgE (1□µg/mL); (ii) Cildir *et al*., anti-hIgE–stimulated hMCs. **(e)** RT-qPCR validation of 18 of the 20 most upregulated DEGs in DS cells (adjusted p□<□0.00001 and |log₂(FC)|□≥□2.5) in naïve, ALL, PN-challenged ALL, and DS cells 2□h after treatment or challenge (n□=□6). Two genes were excluded due to insufficient primer specificity. Red arrows indicate genes significantly different between PN-challenged ALL and DS cells. Data are presented as box plots with median and interquartile range. For normally-distributed groups, one-way ANOVA followed by Bonferroni multiple correction test (*vs.* DS) was applied. Otherwise (*MAFA*, *CHST1*, *GZMB*, *TMEM151A*, *PXDC1*, *PRRX2*), Kruskal-Wallis test followed by Dunn’s multiple correction test was used. *p□<□0.05; **p□<□0.01; ***p□<□0.001; ****p□<□0.0001. PC = principal component.

To determine whether this transcriptional profile was specific to desensitization, we compared the set of upregulated genes in DS cells with publicly available RNA-seq datasets of allergen-challenged hMCs^38, 39^. Only 5% of the total upregulated genes overlapped across the three datasets, with 81 out of 124 genes being uniquely enriched in DS cells (**Fig. 4d**), supporting the specificity of the desensitization-associated program. To validate these findings, we selected the 20 most upregulated genes in DS cells (adjusted *p* < 0.0001 and |log_2_FC| ≥ 2.5) and assessed their expression by RT-qPCR in naïve, ALL, PN-challenged ALL and DS cells 2 h after treatment or challenge. Due to insufficient primer specificity, two genes were excluded, and 18 analyzed. Of these, 12 genes (66.7%) were confirmed to be upregulated in DS cells compared to ALL controls, and 7 (red arrows; ∼40% of the total 18 genes) were significantly elevated relative to PN-challenged ALL cells (**Fig. 4e**). These data validate that desensitization induces a distinct and reproducible transcriptional signature in MCs.

### Desensitization imprints an immunoregulatory transcriptional profile in MCs

To better understand the functional implications of the transcriptional program induced by desensitization, we conducted gene set enrichment analysis (GSEA) across all detected genes from RNA-seq. Using gene ontology (GO) annotations, we identified a broad enrichment of upregulated biological processes in DS cells, primarily related to immune regulation, including “positive regulation of leukocyte cell-cell adhesion”, “regulation of T cell activation” or “chemokine receptor binding” (**Fig. 5a**). Consistent results were obtained when functional annotation analysis was performed on the 168 upregulated DEGs alone (**Fig. 5b**). GSEA based on the hallmark gene dataset further revealed 13 significantly enriched pathways related to immune regulation in DS cells, including “IL-2-STAT5 signaling”, “inflammatory response”, “p53 pathway” or “TNFA signaling via NFκB” (**Fig. 5c**). Moreover, regulatory target gene dataset demonstrated enrichment for transcriptional programs associated with EGR2, NFκB and BACH2 transcription factors (**Fig. S5b**). Together, these analyses suggest that desensitization imprints an immunoregulatory transcriptional profile in MCs. Notably, enrichment of “IL-2-STAT5 signaling” and “regulation of T cell activation” terms is consistent with a previous report indicating that DS MCs promote FoxP3^+^ regulatory T-cell expansion from naïve CD4^+^ T-cells^27^. However, the impact of DS MCs on CD4^+^ T-cell dynamics within an allergic context remains unknown. To address this question, we modeled an allergic environment by isolating splenocytes from PN-allergic mice and co-culturing them with naïve, ALL and DS BMMCs, generated following a polyclonal desensitization protocol^33^. Proliferation of memory/activated CD4^+^CD44^+^ T-cells was assessed by flow cytometry three days after PN challenge (within a four-day co-culture period) (**Fig. 5d**). Although co-culture with BMMCs modestly increased basal proliferation irrespective of condition, allergen challenge specifically enhanced proliferation of CD4^+^CD44^+^ T-cells in the presence of DS BMMCs compared with naïve or ALL BMMCs, or in the absence of BMMCs (**Fig. 5e**). Taken together, these results indicate that DS MCs specifically regulate allergen-specific CD4^+^ T-cell proliferation upon allergen provocations in allergic conditions.

**Figure 5.**
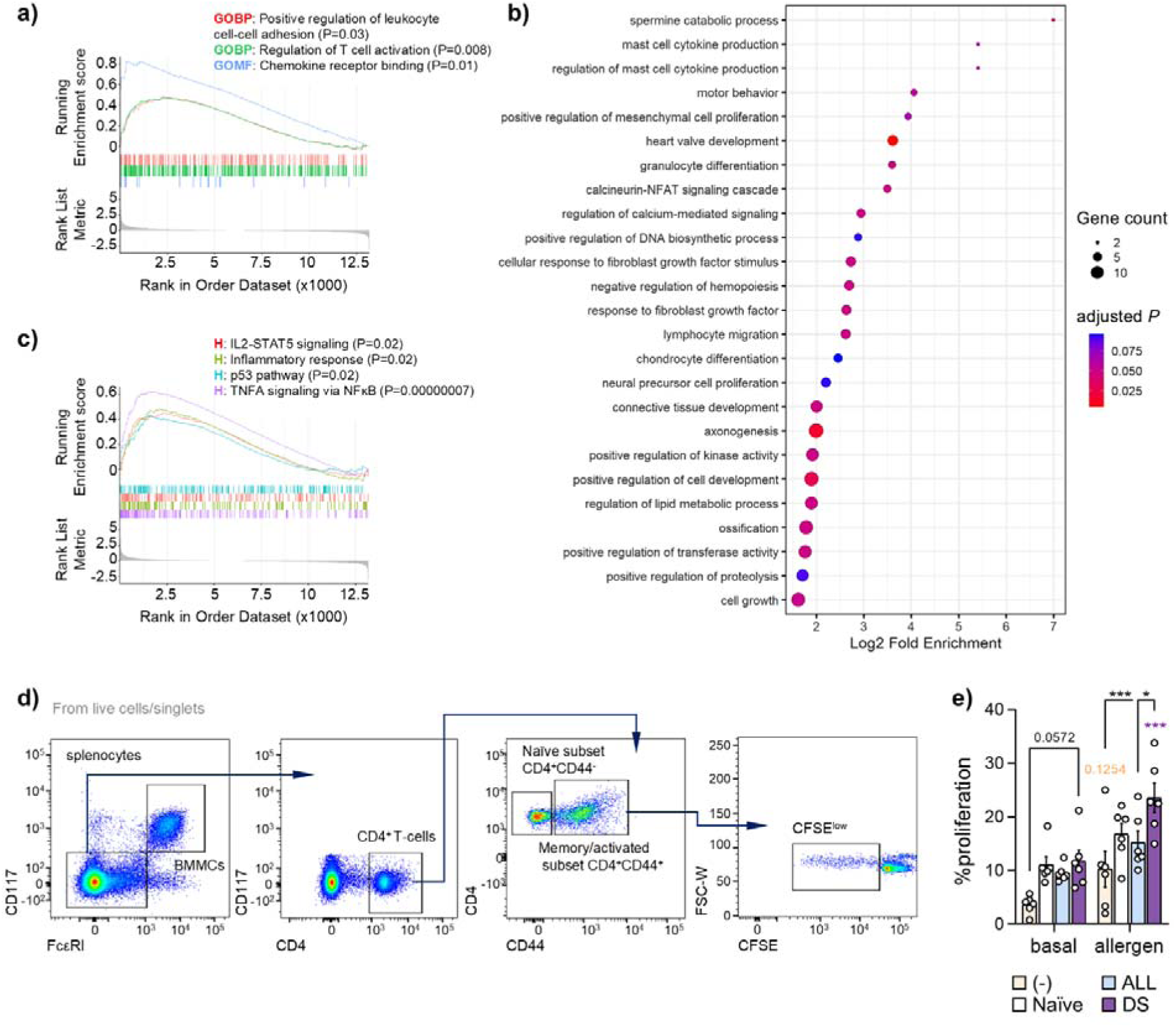
Immunoregulatory features of desensitized mast cells. **(a)** Gene set enrichment analysis (GSEA) of desensitized (DS) cells using Gene Ontology (GO) collection of terms across all expressed genes (n□=□13,168), ranked by log₂ fold change *vs.* allergic (ALL) cells. **(b)** Dot plot of GO enrichment analysis. Dot size represents the number of genes associated with each GO term, and dot color indicates adjusted p-value. **(c)** GSEA of DS cells using Hallmark collection of terms. **(d–e)** CFSE-labeled splenocytes from peanut (PN)-allergic mice were co-cultured with naïve, ALL, or DS bone marrow–derived mast cells (BMMCs). BMMCs were sensitized with pooled sera from PN-allergic mice. Proliferation of memory/activated CD4^⁺^CD44^⁺^ T-cells (CFSE^low^ gate) in monocultures or co-cultures with BMMCs under basal conditions or following PN stimulation; (n□=□6). Data are presented as mean ± SEM. Statistical comparisons were performed using two-way ANOVA followed by Bonferroni multiple comparison test. Colored asterisks indicate significant differences relative to matched basal controls. *p□<□0.05; ***p□<□0.001. CFSE = carboxyfluorescein succinimidyl ester.

## DISCUSSION

MC desensitization is a critical determinant of the safety and efficacy of AIT, which is the only allergen-specific treatment that holds curative potential^18^. However, the mechanisms that operate during MC desensitization are incompletely defined and even controversial^21^. Here, using a polyclonal platform, we demonstrate that MC desensitization is not merely a state of passive hyporesponsiveness, but rather involves coordinated remodeling of IgE/FcεRI signaling and transcriptional programs. We show that desensitization induces time-dependent, allergen-specific IgE internalization while preserving proximal signaling activity yet selectively limiting downstream propagation upon subsequent allergen challenge. Despite largely preserved mitochondrial respiration, desensitized MCs exhibit reduced allergen-induced glycolytic responses and acquire a distinct transcriptional program enriched in immunoregulatory pathways. Together, these findings support a model in which desensitization entails selective signal propagation and cellular reprogramming, potentially contributing to immune modulation and long-term tolerance during AIT.

In contrast to its circulating analogue, the basophil, MCs are tissue-resident long-lived and can retain IgE on the surface for months in the absence of ongoing IgE production^40, 41^, which place them as critical players in IgE-mediated allergic reactions^42^. Intriguingly, during the build-up phase of AIT, despite rising IgE levels, gradual exposure to increasing allergen doses renders MCs hyporesponsive^4, 12^. Although several studies have sought to elucidate the molecular basis of MC desensitization, conflicting findings, particularly regarding IgE/FcεRI internalization, have limited mechanistic consensus. Notably, most prior investigations relied on monoclonal IgE systems in murine models^27, 28, 29, 30, 34^, which may not fully recapitulate the complexity of the polyclonal IgE repertoire present in human allergic individuals^31^. To address this limitation, we employed a human polyclonal desensitization protocol^33^ designed to better reflect physiological IgE diversity. Using this platform, we demonstrate that polyclonal desensitization robustly attenuates allergen-induced degranulation across allergens of distinct origin. To our knowledge, this approach provides the first non–allergen-restricted framework for investigating desensitization mechanisms in human MCs.

Our findings support a model in which desensitization induces allergen-specific IgE internalization while preserving the integrity of the degranulation machinery. By directly comparing monoclonal and polyclonal strategies, we demonstrate that IgE internalization is time-dependent: it is partial immediately after desensitization, whereas 72Lh later allergen-specific IgE is no longer detectable at the cell surface, and allergen-driven degranulation is abrogated. Notably, residual IgE 72Lh post desensitization remained responsive to anti-IgE stimulation, indicating the retention of non-allergen-specific IgEs. Together, these observations suggest that allergen-specific hyporesponsiveness in MCs cannot be solely attributed to a total depletion of allergen-specific IgE. Previous reports have shown IgE/FcεRI reorganization at the cell surface following low-dose allergen exposure^43^, thus it is conceivable that desensitization promotes spatial rearrangement or compartmentalization of IgE/FcεRI complexes, thereby limiting the formation of large receptor clusters required for efficient signal propagation and degranulation. Such a model would reconcile preserved proximal signaling with impaired downstream effector responses observed in our system.

Our data further indicate that IgE/FcεRI signaling is not completely abrogated during desensitization. We observed phosphorylation of SHIP-1, a phosphatase known to negatively regulate FcεRI-dependent degranulation by counteracting proximal signaling events downstream of Syk^44^. The increased SHIP-1 phosphorylation during desensitization may therefore contribute to dampening effector responses without fully extinguishing upstream signaling. In parallel, we detected phosphorylation of LAT, a key adaptor protein required for efficient signalosome assembly following IgE/FcεRI engagement^45^. However, despite some LAT activation during desensitization, subsequent allergen challenge failed to propagate phosphorylation to downstream effectors such as PLCγ, Akt, or ERK in desensitized MCs. These findings suggest that desensitization imposes a threshold constraint on LAT-dependent signaling, resulting in partial uncoupling between proximal activation and distal effector pathways, potentially secondary to impaired allergen recognition. Although the functional role of LAT Y200 remains incompletely characterized, recent evidence indicates that proper spatial organization of LAT is necessary for downstream signal transduction^46^. Whether Y200 phosphorylation specifically modulates LAT-dependent signaling in MCs warrants further investigation.

Desensitization preserved elements of proximal IgE/FcεRI signaling, which prompted us to examine whether this state was accompanied by broader cellular reprogramming. To our knowledge, this is the first study to assess metabolic and transcriptomic adaptations in MCs following desensitization. Untargeted metabolomics and Seahorse analyses revealed that desensitization does not induce major global alterations in mitochondrial respiration. Minor differences observed in acute OCR and spare respiratory capacity were consistent with reduced allergen recognition rather than intrinsic mitochondrial dysfunction. The absence of allergen-specific degranulation in desensitized MCs, likely resulting from the lack of allergen-IgE recognition, was accompanied by a dampening in glycolytic activity (ECAR), indicating that metabolic quiescence parallels functional hyporesponsiveness.

On the other hand, transcriptomic profiling revealed that desensitization induces a distinct gene expression program in MCs. Comparison with previously published RNA-seq datasets from single-challenged hMCs^38, 39^, together with validation of the most upregulated transcripts, confirmed that this signature is specific to the desensitized state rather than a consequence of acute activation. Among the genes selectively enriched in desensitized MCs, several candidates may provide mechanistic insight into this phenotype. Notably, *NOTUM*, a secreted antagonist of Wnt signaling^47^, was among the top upregulated transcripts. Wnt-3a has been associated with type 2 inflammatory signatures in asthma^48^ and enhanced expression of pro-inflammatory cytokines such as IL-8 and CCL-8^49^. Interestingly, IL-8 has been recently reported to be reduced in DS MCs^50^. Therefore, increased *NOTUM* expression may reflect a shift toward attenuation of pro-inflammatory signaling pathways. Additional candidates include *RRAD*, *IL27RA*, *CHST1*, and *XIRP1*: *RRAD* encodes a Ras-related regulator implicated in the restriction of glycolysis through modulation of GLUT1 trafficking^51^, raising the possibility that its upregulation contributes to the reduced glycolytic ATP production observed in DS MCs; IL-27 serum levels increased in allergic patients following AIT^52^ and its alpha subunit receptor (IL27RA) has been reported to mediate inhibitory signals limiting MC degranulation^53^; CHST1-dependent sulfation pathways have been linked to regulation of leukocyte infiltration and inhibitory lectin signaling^54, 55^; and *XIRP1*, an actin-binding protein involved in cytoskeletal organization^56^, may further influence degranulation dynamics. Collectively, these transcriptional changes suggest that desensitization may engage multiple regulatory modules (metabolic, cytokine-mediated, and cytoskeletal) that converge to fine-tune MC responsiveness.

GSEA and GO analyses of both the full transcriptome and the upregulated gene subset revealed broad enrichment of immune-related functions in desensitized MCs. Given the partial overlap between activation– and desensitization-associated transcripts, the enrichment of pathways such as “TNFA signaling via NFκB”^57^ and “inflammatory response” is not unexpected. However, increasing evidence supports a regulatory dimension of MC biology. MC-derived IL-2 production^58, 59^, p53 signaling^60, 61^, and transcriptional programs involving BACH2^62^ have been implicated in restraining allergic inflammation. In addition, the tolerogenic properties of MCs and their capacity for intercellular communication have recently been highlighted^27^. In this context, our functional data indicate that desensitized MCs can modulate allergen-driven proliferation of CD4^⁺^CD44^+^ T-cells, supporting the notion that desensitization imprints an immunomodulatory state in MCs.

In summary, we demonstrate that desensitization induces time-dependent internalization of allergen-specific IgE, thereby limiting subsequent allergen recognition and downstream degranulation-associated responses, including signaling transduction and glycolytic activation. Concurrently, incremental allergen exposure during desensitization preserves selective IgE/FcεRI signaling sufficient to drive a distinct transcriptional program. Together, these findings support a model in which MC desensitization entails selective signaling and transcriptional reprogramming, potentially contributing to immune modulation and long-term tolerance during AIT.

## METHODS

### Patients

This study was approved by the Institutional Research Ethics Committee (approval numbers 4460 and R-0074/23). Patients were recruited from the following Spanish centers: Hospital Universitario de La Princesa, Hospital Infantil Universitario Niño Jesús, and Clínica de Asma y Alergia Dres. Ojeda. Written informed consent was obtained from all participants prior to inclusion. Serum samples were aliquoted and stored at –80°C until use. Clinical characteristics from each patient were registered, and allergen-specific IgE levels were assessed by ImmunoCAP (Phadia, Thermo Fisher) or ALEX (Allergy Xplorer, Macroarray Diagnostic). A detailed description of these data can be found in **Table S1**.

### Animals

Eight-to-twelve-week-old female mice were obtained from Charles River Laboratories Inc. All mice were maintained on a 12-h light-dark cycle and fed an autoclaved chow diet and sterile water *ad libitum*. All procedures were approved by the Environmental Council of the Community of Madrid with PROEX references 45.2/20 and 9.7/23. Ethical regulations for animal research were strictly followed.

### LAD2 culture

The LAD2 cell line was kindly provided by Drs. Metcalfe and Kirshenbaum (National Institute of Allergy and Infectious Diseases, NIAID, USA)^63^. Cells were cultured in StemPro-34 medium containing 2.5% nutrient supplement, 1% GlutaMAX (Thermo Scientific), 1% penicillin/streptomycin (Biowest) and 100 ng/mL recombinant human stem cell factor (rh-SCF; R&D Systems). Cell media was changed weekly, and cellularity was maintained below 0.5×10^6^ cells/mL. Before all experiments, LAD2 cells were pre-treated with 10 ng/mL rh-IL-4 (R&D Systems) for four to five days to enhance FcεRI expression^64^.

### BMMC culture

Bone marrow from femur and tibia of non-allergic C57BL/6 female mice was used to generate BMMCs as previously described^65^. Erythrocyte-free bone marrow cells were cultured for four weeks in Iscoves’s Modified Dulbecco’s medium (IMDM; Gibco) supplemented with 10% fetal bovine serum (FBS; Cytiva), 1% minimal essential medium (Thermo Scientific), 1% sodium pyruvate (Biowest), 1% penicillin/streptomycin, 1% non-essential amino acids (Biowest), 10 ng/mL recombinant murine (rm)-SCF (PeproTech) and 5 ng/mL rm-IL-3 (PeproTech). Cell media was changed weekly, and cellularity was kept below 0.5×10^6^ cells/mL. BMMC differentiation was assessed by flow cytometry. Mature BMMCs (>90% CD117^+^/FcLRI^+^) were obtained and used for functional experiments in weeks four and five of culture.

### Primary hMC culture

Primary hMCs were generated from circulating CD34^+^ hematopoietic progenitors purified from buffy coats of healthy donors provided by the Transfusion Center of the Community of Madrid. First, human peripheral blood mononuclear cells (PBMCs) were isolated from whole blood using density gradient separation (Pancoll human; PAN Biotech). CD34^+^ hematopoietic progenitors were isolated from 5×10^7-108^ PBMCs using the EasySep Human CD34 Positive Selection Kit II (STEMCELL Technologies). To do that, PBMCs were suspended in 1 mL of EasySep buffer (STEMCELL Technologies) and incubated for 10 min at room temperature (RT) with 100 µL magnetic anti-CD34 antibody cocktail. Subsequently, 27 µL of RapidSpheres were added and incubated 5 min at RT. Cells were then placed into an EasySep Magnet (STEMCELL Technologies) and incubated for 3 min before pouring off the supernatant. After 5 x 3-min washes, CD34^+^ cells were obtained for culture.

For the first two weeks, CD34^+^ progenitors were cultured at 0.5×10^6^ cell/mL in supplemented StemPro-34 adding 100 ng/mL rh-IL-3 and 50 ng/mL rh-IL-6 (STEMCELL Technologies). For the third and fourth weeks, only rh-IL-6 supplemented StemPro-34 fresh media was added. From week five to eight, supplemented media without any additional interleukins was used. Medium was renewed every 3-4 days and mature hMCs were used for functional assays from week five onwards when the culture reached a differentiation >30% of CD117^+^/FcεRI^+^ cells, assessed by flow cytometry.

### LAD2 and hMC desensitization and activation experiments

LAD2 and hMCs (10^6^ cells/mL) were sensitized overnight with sera from patients allergic to PN, egg white, milk, grass pollen and cat dander with specific IgE levels > 20 kU/L at selected concentrations (**Table S1**). Sera dilution was chosen based on a maximal LAD2 degranulating response of 20-40%. Allergen desensitization was performed the next day following a polyclonal protocol previously described^33^. For this purpose, ALL cells were placed in a flat-bottom well plate at a cell density of 10^6^ cells/mL and desensitized by adding increasing doses of the specific allergen in 10 min intervals: PN extract (#XPF171D3A25, Stallergenes Greer), milk powder (Hacendado) egg white (#E0500, Sigma-Aldrich), grass pollen (#40.10.85.10, CITEQ biologics) or cat dander (#10.01.51.10, CITEQ biologics).

For MC activation, ALL and DS cells were challenged during 10 min with the specific allergens at 20 μg/mL. Activation was stopped at 4°C before analyzing MC responses by flow cytometry (CD63 and CD107a expression in CD117^+^FcεRI^+^ gated population) and β-hexosaminidase assay^33^. Cells stimulated with media or goat polyclonal anti-hIgE (20 μg/mL; #I6284, Sigma or #ab9159, Abcam) were used as negative and positive controls of degranulation, respectively. Additionally, naïve cells were used as negative control for sensitization. In addition, IgE MFI was evaluated in ALL and DS cells by flow cytometry within the CD117^+^/FcεRI^+^ gated population while FcLRI MFI was measured within the CD117^+^ gated population.

### Kinetics of IgE internalization

IgE surface levels were assessed at 0, 24, 48, and 72Lh using two desensitization strategies: monoclonal and polyclonal. For monoclonal desensitization, BMMCs were sensitized with 5 µg/mL OVA-specific monoclonal IgE (clone EC1; Chondrex Inc.) and desensitized according to the protocol described by Oka *et al.*^28^ Culture medium was refreshed 1–2 days prior to sensitization. For polyclonal desensitization, LAD2 cells were sensitized and desensitized as described above. MC degranulation was evaluated 72Lh post-desensitization by stimulation with OVA (10 µg/mL; Sigma-Aldrich) or anti-mIgE (10 µg/mL; Thermo Scientific) in BMMCs, and with allergen (20 µg/mL) or anti-hIgE (20 µg/mL) in LAD2 cells. IgE MFI was quantified by flow cytometry as described above. Naïve and unstained cells were used as negative and staining controls, respectively.

### Mouse splenocyte isolation and BMMC co-culture

Splenocytes were obtained from PN-allergic C57BL/6 female mice. Mice were intra-gastric sensitized by administrating 4 mg of PN butter (∼1 mg of protein; Capitán Mani) and 5 µg of cholera toxin (Sigma-Aldrich) in 200-500 µL of PBS by oral gavage once a week for 4 weeks^40, 66^. Following euthanasia, spleen was collected and splenocytes were isolated as previously described^67, 68^. For CFSE staining, splenocytes were suspended at 8×10^6^ cells/mL in RPMI media (Gibco) containing 1% GlutaMAX and 25 mM HEPES, and supplemented with 10% FBS, 1% penicillin/streptomycin, 1 % sodium pyruvate, 1% non-essential amino acids and 55 µM β-mercaptoethanol. Then, 4.3 µL of CFSE at 2.79 mg/mL in DMSO (Sigma) were added and incubated for 5 min at RT protected from light. Afterwards, 3 x 5-min washes (406g at 4°C) were performed. CFSE-stained cells were suspended in complete RPMI and 0.4×10^6^ cells per condition were used^40^.

For co-culture experiments, mature BMMCs were sensitized with sera from PN-allergic mice at 1:100 dilution (mice pool sera A [PN-specific IgE = 160.57 ng/mL]; mice pool sera B [PN-specific IgE = 10-39 ng/mL]), and desensitized (10^6^ cells/mL) performing a 9-step polyclonal desensitization protocol, as described^33^. Subsequently, 0.4×10^6^ cells were suspended in complete RPMI and added to CFSE-stained spleen cells (ratio 1:1). Then, 25 µg/mL of PN extract were added 24 h after. Co-cultures were incubated for 3 additional days. T-cell proliferation within CD4^+^ T-cell subpopulations was assessed by measuring CFSE^low^ population by flow cytometry.

### Flow cytometry

In all experiments, cells were suspended in ice cold FACS buffer (0.5% bovine serum albumin [BSA; Sigma-Aldrich] and 2.5 mM EDTA in PBS). Fc-blocking was performed for 15 min with 1:50 of Human TruStain FcX (Biolegend) for LAD2 and hMCs, or anti-mouse CD16/CD32 (BioLegend) for BMMCs. Then, cells were incubated with a combination of fluorophore labelled antibodies (**Table S2**) on ice for 30 min protected from light. Cell viability was assessed with eFluor-780 dye 1:4,000 (eBioscience). After staining, cells were washed, suspended in 150 μL of FACS buffer and analyzed on a FACSCantoII or FACSLyric flow cytometer flow cytometers (Becton Dickinson). On average, 10,000 live and single cells were recorded. For co-culture experiments, 200,000 events were acquired. Data were analyzed using *FlowJo v10* software (Beckton Dickinson). Fluorescence minus one (FMO) controls were used for gating strategy.

### β-hexosaminidase enzymatic activity

MC supernatants were used for β-hexosaminidase assays^69^. Briefly, 45 μL of the supernatant plus 45 μL of the β-hexosaminidase substrate solution (2 mM p-nitrophenyl N-acetyl b-D-glucosamine in 100 mM citrate buffer pH 4.5; #N9376, Sigma Aldrich) were incubated per duplicate into a flat-bottom 96-well plate and incubated at 37°C protected from light for 45 min. The reaction was stopped by adding 90 μL of 1M NaOH. In addition, 45 μL of culture media alone were used to determine the absorbance background, and 45 μL of TritonX100-lysed cells were employed to measure total levels of β-hexosaminidase within cells. Optical density (OD) was measured at 405 nm with a microplate reader (GloMAX Discover; Promega). Percentage of β-hexosaminidase activity was calculated using the following formula: % of degranulation = [(OD supernatant – background)/(OD total lysis – background)]x100.

### IgE cell surface stripping

IgE stripping from LAD2 cell membrane was performed using a mild acetic acid wash, as previously reported^70^. Briefly, 10^6^ cells/mL ALL and DS cells were incubated in 50 μL ice-cold acetic acid buffer pH 4 (50 mM acetate, 85 mM NaCl, 10 mM EDTA and 0.03% BSA) for 5 min. Then, an equal volume of supplemented media culture was added, and cells were centrifuged at 265 g for 5 min at 4°C to collect the IgE contained in cell supernatant. Afterwards, the stripped-IgE (ALL and DS) was used for sensitizing naïve LAD2 cells. Total IgE stripping was confirmed by flow cytometry. In addition, stripped ALL and DS cells were re-sensitized with the same previous allergic serum. The next day, cells were washed and challenged with the specific allergen extracts (20 μg/mL) or anti-hIgE (20 μg/mL) for 10 min, as described above. MC activation was analyzed by flow cytometry and β-hexosaminidase assay.

### Seahorse metabolic flux assays

Analysis of OCR and ECAR rates were performed using the Seahorse XFe96 system (Agilent Technologies). For this purpose, PN-ALL and DS LAD2 cells were seeded in a 96-well seahorse plate pre-coated with 100 μg/mL of poly-L-lysine (Sigma Aldrich) following Agilent recommendations for non-adherent cells. Briefly, ALL and DS cells were centrifuged at 265g for 5 min at RT and suspended in Dulbecco’s Modified Eagle media (DMEM) pH 7.4, supplemented with 5 mM glucose, 2 mM glutamine and 1 mM sodium pyruvate at a cellularity of 1.4-1.6 x 10^6^ cells/mL. Next, 50 µL per condition (*i.e.*, 7-8 x 10^4^ cells) were plated per quintuplicate and centrifuged at 200g for 2 min with low brake at RT. Cells were then incubated at 37°C in a non-CO_2_ incubator for 20 min before adding 130 µL of seahorse media to achieve a final volume of 180 µL. Finally, cells were allowed to rest for 15 additional min as aforementioned before metabolic measures were taken. Basal state was analyzed prior sequential additions of 1.5 μM oligomycin A (Sigma or Agilent Technologies), 1.5 μM FCCP (Sigma) and 1 μM rotenone (Sigma or Agilent Technologies) plus 1 μM antimycin A (Sigma or Agilent Technologies), following the Seahorse XF96 Analyzer protocol specifications. In stimulated conditions, cells were activated with PN extract (20 μg/mL) or anti-hIgE (10 μg/mL) before oligomycin injection. To determine the differences in ATP rates, 150 µL of media was added to achieve a final volume of 200 µL and 2.5 μM BAM15 (Agilent Technologies) was used instead of FCCP. A total of three measurements of OCR and ECAR were taken for each drug injection. For well-to-well normalization, cell nuclei were Hoechst-stained (Hoechst 33342; Thermo Scientific) and counted using a BioTek Cytation 5 Cell Imaging Multimode Reader (four regions per well). Analyses were conducted using Agilent *Seahorse Analytics* web (https://seahorseanalytics.agilent.com).

### Metabolomic analysis

For metabolomic analysis, LAD2 cells were sensitized with 100 ng/mL myeloma-derived hIgE (#401152, EMD Milipore) and desensitized (0.5×10^6^ cells/mL) following a polyclonal anti-IgE protocol^33^. Supernatants from ALL or DS cells were collected after desensitization and stored at –80°C. Data acquisition and analysis was performed at CEMBIO as previously described^71, 72, 73^. Briefly, 50 μL of supernatant from each condition were separated and put together with 10 μL of an internal standard mix. Samples were then deproteinized by centrifugation at 2,000g for 80 min at 4°C on a Centrifree ultra-centrifugation device with a cut-off porous of 30 kDa (Millipore Ireland Ltd.). Additional quality controls were prepared by pooling equal volumes of all samples to provide evidence about the stability, performance and reproducibility of the analytical technique. Sample analysis was conducted following two different methodologies to study cationic or anionic metabolites using capillary electrophoresis (7100, Agilent Technologies), operated in normal or reverse polarity, coupled to a time of flight-mass spectrometry in positive (6230) or negative (6224, Agilent Technologies) ionization mode, respectively. Data processing was performed using Agilent *MassHunter Profinder* software (B.10.0.2).

For metabolite identification, chemical signals were annotated by monosotopic mass and relative migration time using an in-house library available at CEUMassMediator (CE-MS search; http://ceumass.eps.uspceu.es/index_cesearch.xhtml)^73^. Unsupervised PCA and heatmap analysis were performed using *R* (version 4.3.3)^74^. PCA on detected metabolites was performed to assess data quality and clustering of samples. This model was log_10_ scaled and evaluated by the first and second principal components. Heatmap analysis was carried out using complete agglomerative hierarchical clustering applied to the Euclidean distance matrix of the expression levels of the detected metabolites.

### Western blot analysis

For signaling pathway analysis during desensitization, lysates of 0.5×10^6^ PN-ALL LAD2 cells were collected at different desensitization steps (step 1 – 0.1 ng/mL; step 3 – 0.313 ng/mL; step 6 – 2.5 ng/mL; step 9 – 10 ng/mL), together with dose-matching ALL cells challenged for 10 min. To study the pathways triggered in ALL and DS cells upon allergen challenge, lysates from PN-ALL and DS cells were obtained 1 min after PN challenge (20 µg/mL).

Total protein lysates were collected using RIPA lysis buffer (1% NP40, 0.5% sodium deoxycholate and 0.1% SDS in distilled H_2_O) containing 1% protease (cOmplete, Roche Diagnostics) and 1% phosphatase inhibitors (PhosSTOP, Roche Diagnostics), reaching a final concentration of 5,000 cells/μL. Subsequently, protein lysates were mixed with 4x Laemmli sample buffer (Bio-Rad) with 10% β-mercaptoethanol, heat-denatured at 100°C for 5 min and resolved in a homemade 10% SDS-acrylamide gel (Bio-Rad). Additionally, PageRuler Plus protein ladder (Thermo Scientific) was added as an indicator of protein size. Electrophoresis was performed at constant voltage (80-120 V) using Tris-Glycine-SDS running buffer (Bio-Rad). Proteins were transferred for 1 h at constant amperage (400 mA) to a 5-min methanol-activated PVDF membrane (Bio-Rad). Membranes were blocked in 3% BSA in Tris-buffered saline (140 mM NaCl, 10 mM Tris-HCl, pH 7.4) and 0.1% Tween20 (TBS-T) for 1 h at RT and incubated overnight at 4°C with primary antibodies diluted in TBS-T with 5% BSA and 0.01% sodium azide (**Table S2**). After four washes in TBS-T for 10 min, membranes were incubated with goat anti-rabbit or anti-mouse horseradish peroxidase (HRP)-conjugated secondary antibody (1:5,000) for 1 h at RT. Membranes were stripped using the Pierce Restore PLUS blot stripping buffer (Thermo Scientific) according to manufacture specifications. ECL Prime Western Blotting Detection Reagent (Cytiva) was used to visualize protein bands and blot chemiluminescence was detected by ImageQuant 800 system (Amersham). Densitometric values were obtained using *ImageJ* software (version 1.8.0)^75^.

### RNA extraction and RT-qPCR

Total RNA from 1×10^6^ naïve, PN-ALL, PN-challenged and DS LAD2 cells was extracted 2 h post-desensitization or challenge using NucleoSpin RNA/Protein kit (Macherey-Nagel), as recommended by the manufacturer. Cells were centrifuged at 265g for 5 min, homogenized in RP1 lysis buffer supplemented with β-mercaptoethanol (1:100) and stored at –80°C before RNA isolation was performed. Quantity and quality of total isolated RNA were evaluated by NanoDrop (ND-100, Thermo Scientific). cDNA reverse transcription was performed in a SimpliAmp thermal cycler (Applied Biosystems) from 500-600 ng of total RNA using the High-Capacity cDNA RT kit (Applied Biosystems) with the following protocol: 25°C 10 min, 37°C 2 h, 85°C 5 min. Gene expression was evaluated with 10 µM specific mRNA primers (**Table S3**) by real-time qPCR using SYBR green as a reporter (GoTaq(R) qPCR Master Mix; Promega). RT-qPCR amplifications were performed in a QuantStudio 5 Real-Time PCR system, 384-well thermal cycler (Applied Biosystems) with the following protocol: 40 cycles (95°C 15 seconds, 60°C 1 min) followed by a 3-step melt curve (95°C 15 seconds, 60°C 1 min, 95°C 15 seconds). Relative gene expression was calculated using the 2^−ΔΔCT^ method^76^ normalizing to two housekeeping genes: ribosomal 18S (*R18S*) and glyceraldehyde-3-phosphate dehydrogenase (*GAPDH*).

### RNA-seq quantification and analysis

Total RNA from four replicates of naïve, PN-ALL and DS LAD2 cells (2 h post-desensitization) was obtained as aforementioned and sequenced at BGI facilities. DNA library quality control and circularization were followed by amplification to make DNA nanoballs, which were finally sequenced on DNBSEQ-G400 (DNBSEQ Technology) platform (150 bp paired-end sequencing, aiming for at least 20 million clean reads per sample).

Adapter and low-quality reads were filtered out from FASTQ files using *SOAPnuke* software by BGI^77^. Clean files were then subjected to an in-house quality control analysis using *FastQC* (version 0.12.1)^78^. All 12 samples passed this second quality control. *Kallisto* (version 0.46.2)^79^ was employed to pseudo-align reads to the reference human genome GRCh38.p14. On average, more than 22 million reads per sample were processed, yielding pseudo-alignment ratios above 86% in all cases (**Table S4**).

Subsequent data analysis was fully carried out using *R* (version 4.3.3)^74^. Gene-level abundance data were filtered to only keep genes with expression levels above 1 count per million (CPM) in at least four samples, and afterwards, trimmed mean of M-values method was used to normalize data among samples using *edgeR* package (version 4.0.16)^80^. PCA on log_2_(CPM) data was carried out to assess unsupervised clustering of samples.

DEGs between ALL *vs.* naïve samples, and DS *vs.* ALL samples, were identified using *limma* package (version 3.58.1)^81^. For this purpose, a weighted linear model accounting for mean-variance relationship was fitted to log_2_(CPM) normalized data, adjusting p-values by means of Benjamini-Hochberg method to control false discovery rate (FDR) and using *eBayes* function for empirical Bayes moderation of standard errors. Genes with both adjusted p-value ≤ 0.05 and |log_2_FC| ≥ 1 were considered differentially expressed. For sample grouping, complete agglomerative hierarchical clustering was applied to the Euclidean distance matrix of significant DEGs log_2_(CPM) expression levels. Finally, for functional analysis, GSEA and GO enrichment were performed relying on *clusterProfiler* (version 4.10.1)^82^ and *msigdbr* (version 25.1.1)^83^ packages. Redundancy of enriched GO terms was removed using *simplify* function from *clusterProfiler* package.

### Statistical analysis

All analysis were performed using *GraphPad Prism v10* (GraphPad Software, USA) and *R* (version 4.3.3)^74^ software. Quantitative variables are represented as mean ± standard error of the mean (SEM). Normal distribution was determined by D’Agostino-Pearson and Shapiro-Wilk tests. If normal distribution could be assumed, parametric test was used. Correlation analyses were conducted using Pearson correlation test. For multiple comparisons of the media between three or more groups, one or two-way analysis of the variance (ANOVA) tests followed by Bonferroni multiple correction tests were used. For two-way ANOVA tests, data that did not follow normal distribution were logarithmically transformed. On the other hand, non-parametric tests were conducted when normal distribution was not followed. Kruskal-Wallis followed by Dunn’s multiple correction tests were conducted to compare the medians of three or more independent groups, whereas Wilcoxon tests were performed to compare the medians between two dependent groups. A p-value of <L0.05 was considered statistically significant.

## DATA AVAILABILITY

The RNA-seq data generated in this study will be openly available at the NCBI Gene Expression Omnibus (GEO) at https://www.ncbi.nlm.nih.gov/geo/. Metabolomic data will be publicly available at the Metabolomics Workbench (https://www.metabolomicsworkbench.org).

## ACKNOWLEDGMENTS

We thank Drs. Metcalfe and Kirshenbaum for providing the LAD2 cell line. We acknowledge the support of the Cell Culture Department of the Centro de Investigaciones Biológicas (CIB) for conducting the initial proof-of-concept Seahorse experiments. We are grateful to Dr. A. Martínez-Caro for providing the Agilent Seahorse XFp T Cell Metabolic Profiling Kit and for insightful discussions on Seahorse data interpretation. We also thank all allergic patients who voluntarily participated in this study.

## FUNDING

RJS’s laboratory is supported by the SEAIC (BECA20A9); by the Instituto de Salud Carlos III (ISCIII; CP20/00043, PI22/00236, and PI25/00477); by the Ministerio de Ciencia, Innovación y Universidades (MICIU) through project CNS2024-154194 (MCIN/AEI/10.13039/501100011033); by the Fundación Domingo Martínez (FDM2025); and by the Weston Foundation (POP24-11435).

CLS is supported by ISCIII through a Río Hortega contract (CM23/0019). ENB is supported by ISCIII through a Sara Borrell contract (CD23/00125) and by IIS-Princesa (PIM-011) and the UAX Foundation (950.690). ARS is supported by grant PEJ-2024-AI/SAL-GL-32172 from the Youth Employment Program of the Comunidad de Madrid. ESM and AJGC are supported by Formación de Profesorado Universitario (FPU) fellowships from MICIU (FPU23/03341 and FPU24/01211, respectively). PRR is supported by the ISCIII (PI24/01770 and INT25/00101).

CLS, ENB, AV, PRR, and RJS are members of RICORS “Red de Enfermedades Inflamatorias (REI)” (ISCIII; RD24/0007/0018 and RD24/0007/0037).

## CONFLICT OF INTEREST

All authors declare no competing interest regarding this work.

## AUTHOR CONTIBUTION

Conceptualization: CLS, ENB, RJS.

Methodology: CLS, ENB, ARS, JSM, AJGC, SAG, MZD, ESM, LMS, MMH, RJS.

Formal analysis: CLS, ENB, JSM. Investigation: CLS, ENB, ARS, SAG.

Resources: AV, DB, EB, CMC, PO, CB, PRR, FSM.

Data curation: CLS, ENB.

Visualization: CLS, ENB, ARS, JSM, AJGC. Supervision: RJS.

Project administration: RJS.

Writing – original draft: CLS, ENB, RJS. Writing – review & editing: All authors.

Funding acquisition: RJS.

## ABBREVIATIONS

AIT: Allergen immunotherapy
ALL: Allergic
ANOVA: Analysis of variance
BMMCs: Bone-marrow derived mast cells
BSA: Bovine serum albumin
DS: Desensitized
DMEM: Dulbecco’s Modified Eagle media
SEM: Standard error of the mean
ECAR: Extracellular acidification rate
FDR: False discovery rate
GSEA: Gene set enrichment analysis
GEO: Gene expression omnibus
GO: Gene ontology
GAPDH: Glyceraldehyde-3-phosphate dehydrogenase
FcLR: High-affinity Fc ε receptor
h: Human
Ig: Immunoglobulin
IMDM: Iscoves’s Modified Dulbecco’s medium
MCs: Mast cells
m: Murine
OCR: Oxygen consumption rate
OD: Optical density
OVA: Ovalbumin
PN: Peanut
PBMCs: Peripheral blood mononuclear cells
RT-qPCR: Real time, quantitative PCR
r: Recombinant
R18S: Ribosomal 18S
reg: Regulatory
RNA-seq: RNA sequencing
SRA: Sequence read archive
TBS: Tris-buffered saline

## Notes

### Competing Interest Statement

The authors have declared no competing interest.

