## Supplementary Figures and Tables for "Mast cell desensitization induces a distinct IgE-dependent transcriptional program associated with immune regulation"

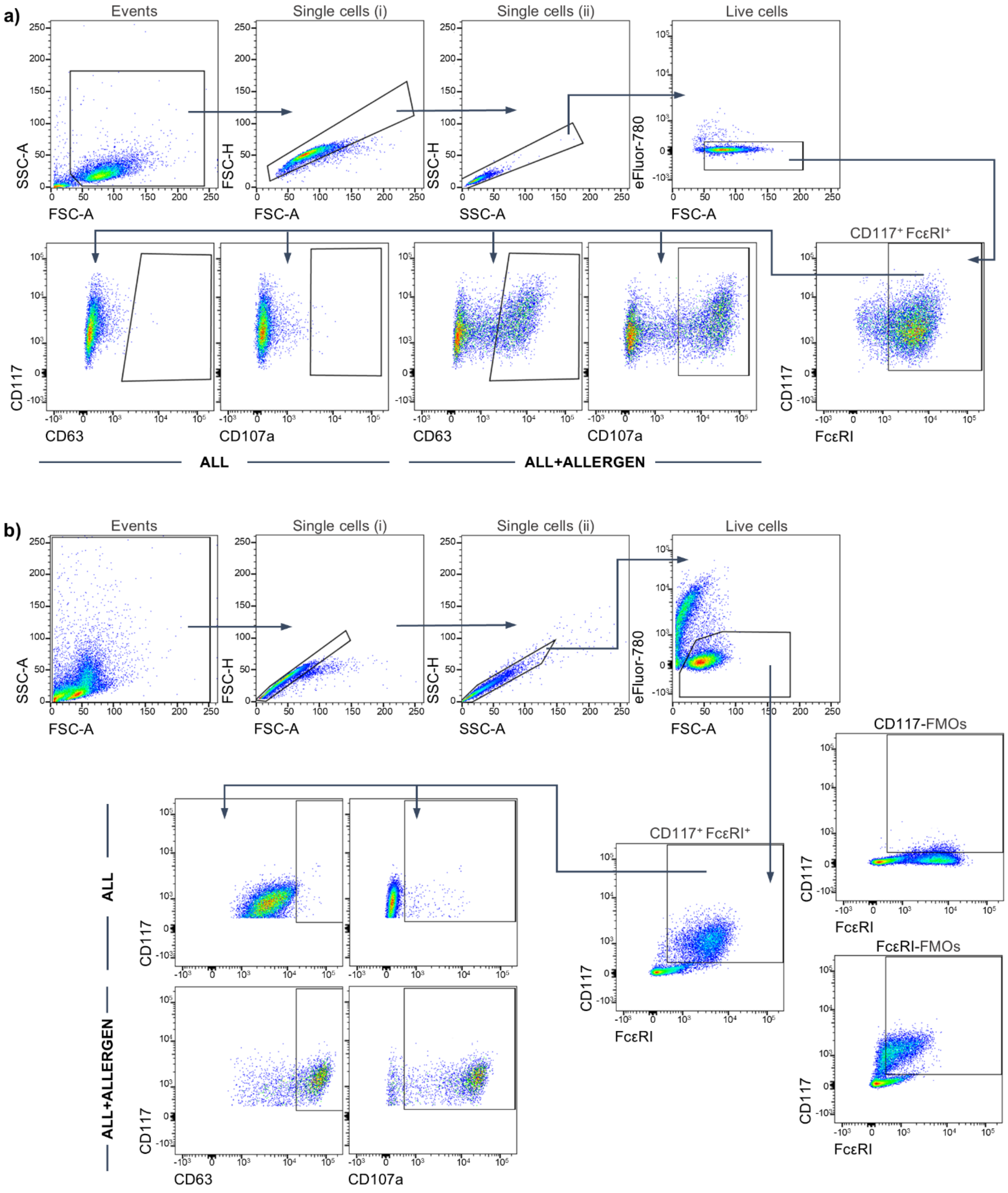


**Supplementary figure 1. Flow cytometry gating strategy for LAD2 and primary human mast cells**. **(a)** Gating strategy for LAD2 cells cultured for 4–5 days in the presence of IL-4. **(b)** Gating strategy for $CD34^+$-derived human mast cells (hMCs). Live cells were identified as eFluor-780–negative events. Doublets were excluded based on forward scatter height *vs.* area (FSC-H *vs.* FSC-A), followed by side scatter height *vs.* area (SSC-H *vs.* SSC-A). hMCs were gated as $CD117^+FcεRI^+$ cells using fluorescence-minus-one (FMO) controls. Activation markers analyzed included CD63 and CD107a. ALL, sensitized mast cells.

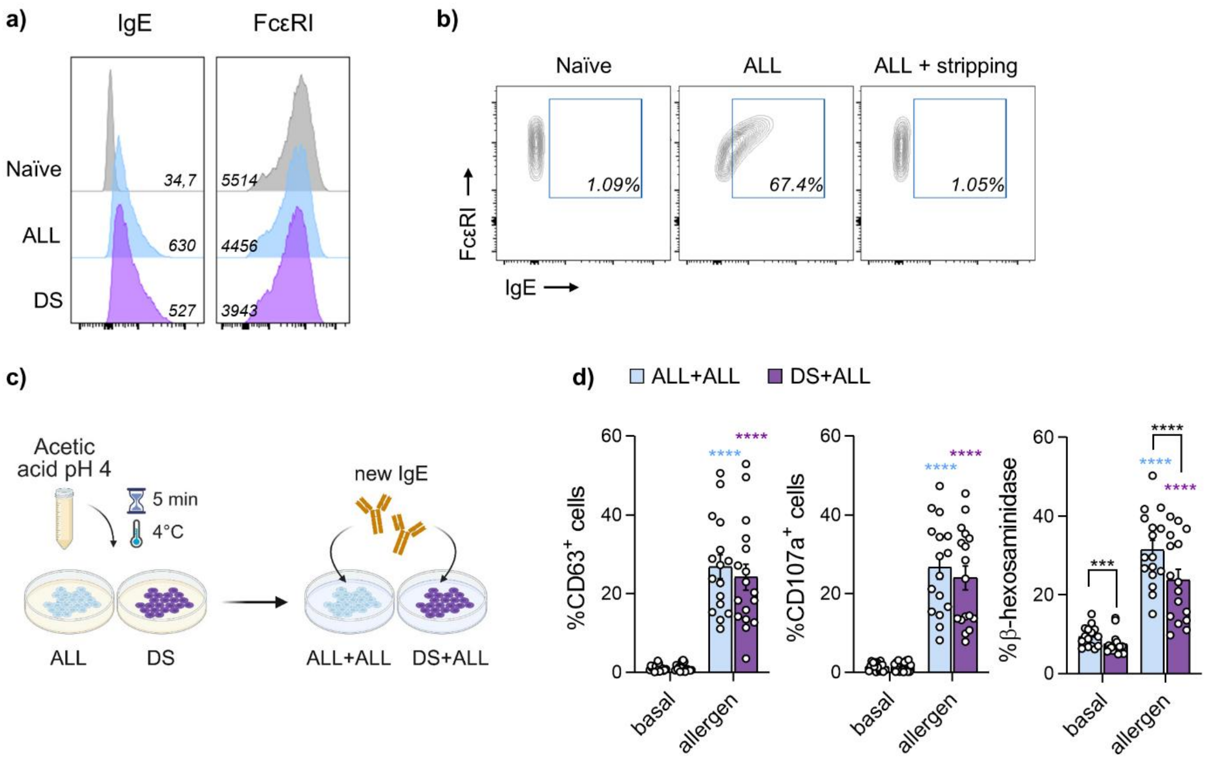


**Supplementary figure 2. IgE expression, stripping efficiency, and functional recovery assays**. **(a)** Representative histograms showing mean fluorescence intensity (MFI) of IgE and FcεRI expression measured by flow cytometry in naïve, sensitized (ALL), and desensitized (DS) LAD2 cells. Cells were sensitized with sera from patients allergic to cat, egg white, peanut, pollen, or milk. **(b)** Representative dot plot showing efficiency of IgE stripping assessed by flow cytometry. **(c)** Experimental design. IgE-stripped ALL and DS LAD2 cells were re-sensitized with the same allergic serum used for initial sensitization (ALL+ALL and DS+ALL conditions, respectively). Image created with BioRender.com. **(d)** CD63 and CD107a surface expression and β-hexosaminidase release following allergen challenge in ALL+ALL and DS+ALL LAD2 cells (n = 17). Data are presented as mean ± SEM. Statistical comparisons were performed using two-way ANOVA followed by Bonferroni multiple comparison test. Data that did not follow normal distribution (CD107a and β-hexosaminidase) was logarithmically transformed before two-way ANOVA was performed. Colored asterisks indicate significant differences relative to matched basal controls. **p < 0.01; ****p < 0.0001.

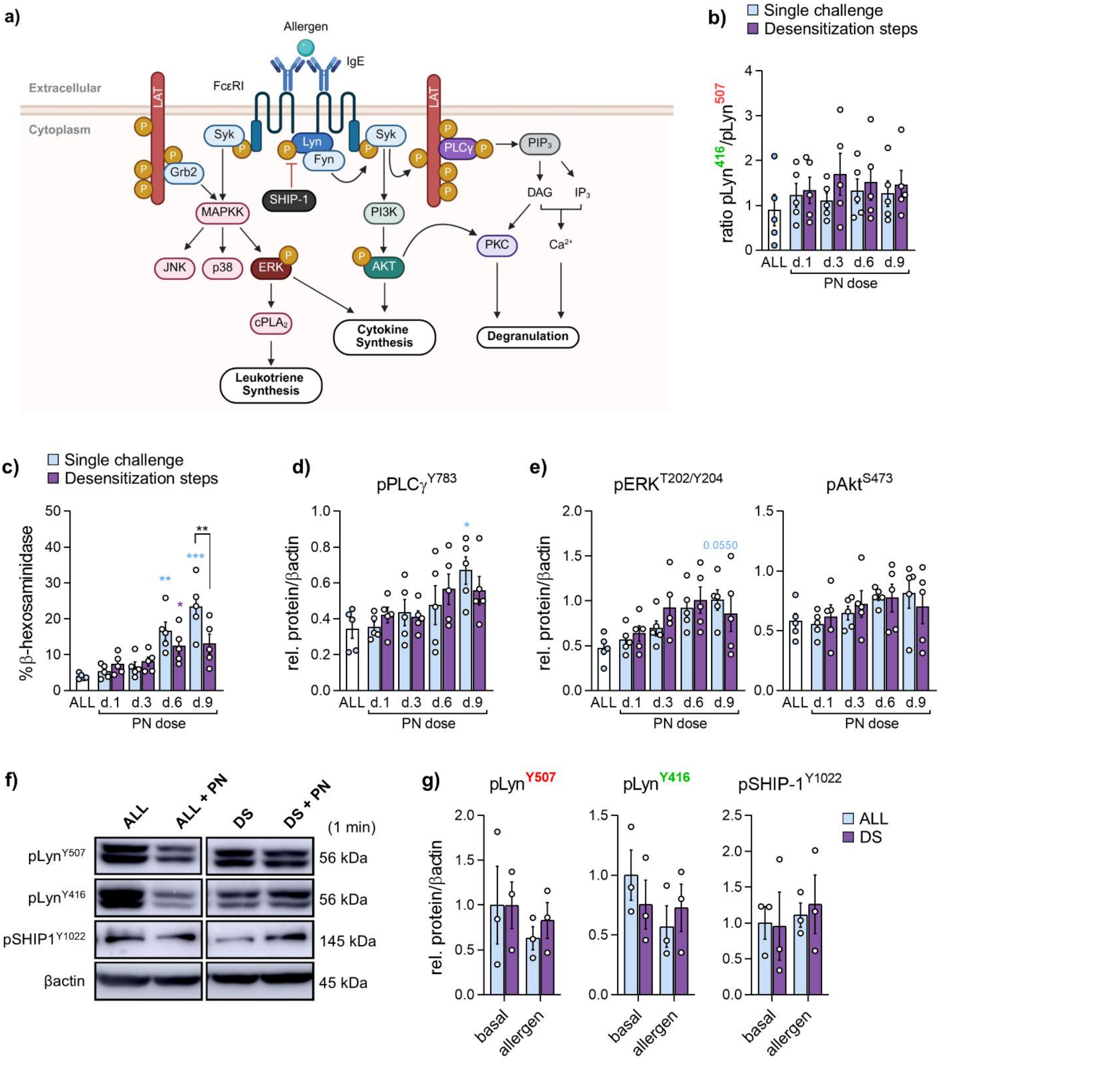


**Supplementary figure 3**. **IgE/FcεRI signaling dynamics during desensitization**. LAD2 cells were sensitized with peanut (PN)-allergic sera and either desensitized or stimulated with single-dose PN challenges matching the cumulative allergen doses reached at different desensitization steps (0.1, 0.31, 2.53, and 10 ng/mL). **(a)** Schematic representation of the IgE/FcεRI signaling cascade. Colored proteins indicate those analyzed by Western blot. Image created with BioRender.com. **(b)** Quantification of the Lyn activating-to-inhibitory phosphorylation ratio (Y416/Y507). **(c)** β-hexosaminidase release during single-dose or cumulative allergen stimulation. **(d, e)** Phosphorylation levels of PLCγ (Y783) **(d)**, ERK (T202/Y204) and Akt (S473) **(e)**, normalized to β-actin; (n = 5). **(f)** Representative Western blot showing Lyn and SHIP-1 phosphorylation in PN-sensitized (ALL) and desensitized (DS) LAD2 cells following 1 min allergen stimulation (20 µg/mL). This Western blot comes from the same gel as Fig. 2f. **(g)** Quantification of Lyn (Y416 and Y507) and SHIP-1 (Y1022) phosphorylation relative to β-actin; (n = 3). Data are presented as mean ± SEM. Statistical comparisons were performed using two-way ANOVA followed by Bonferroni multiple comparison test. Colored asterisks indicate significant differences relative to matched basal controls. $^{*}p < 0.05$; $^{**}p < 0.01$; $^{***}p < 0.001$.

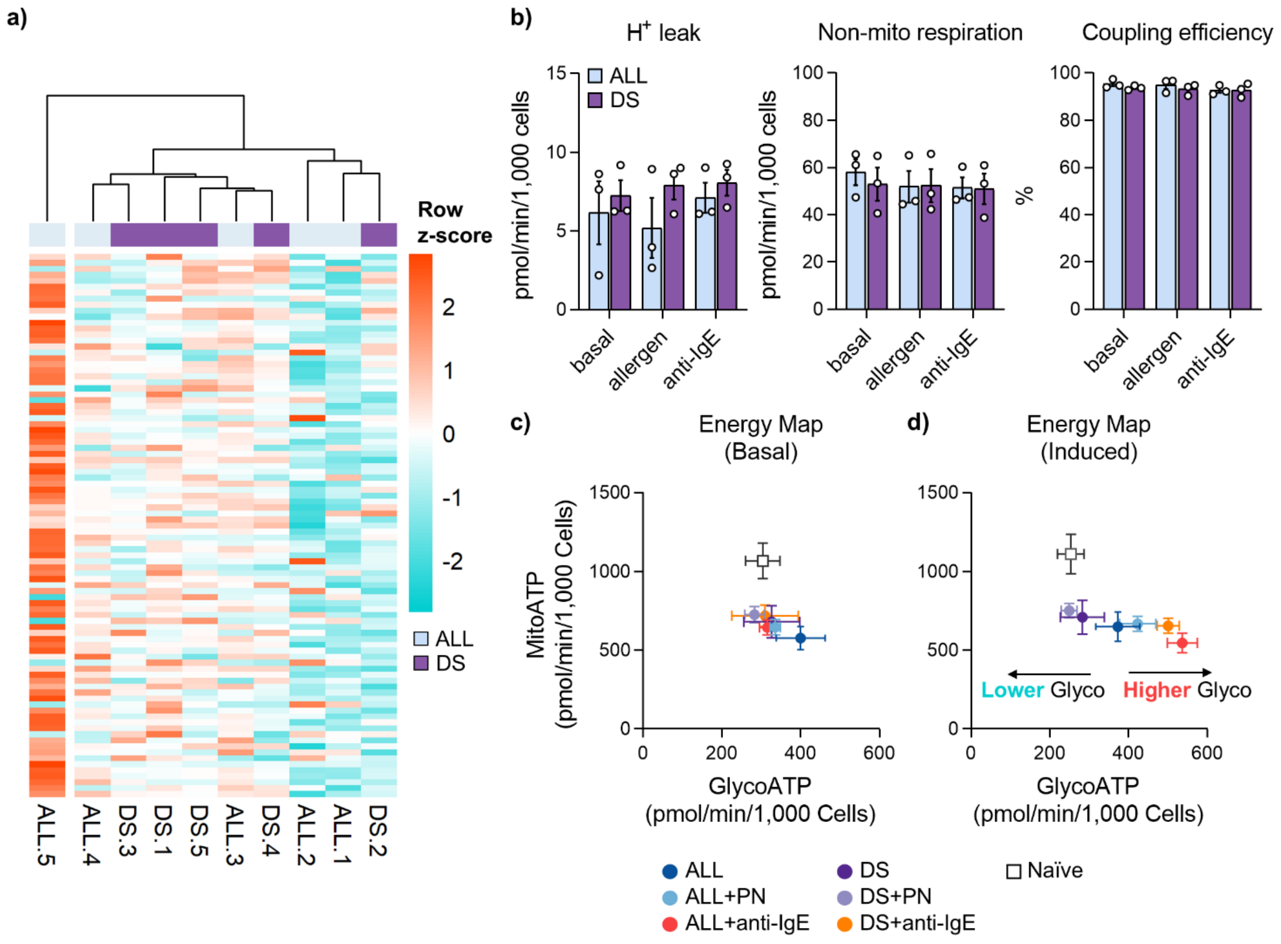


**Supplementary figure 4**. **Metabolic profiling of sensitized and desensitized mast cells. (a)** Heatmap showing hierarchical clustering based on the 91 metabolites identified in sensitized (ALL) and desensitized (DS) LAD2 cells (n = 5). ALL cells were sensitized with human (h) myeloma IgE (100 µg/mL) and desensitized using a 7-step anti-hIgE protocol. Rows are scaled by metabolite abundance. Blue, downregulated; red, upregulated. **(b)** Quantification of OCR-derived metabolic parameters in peanut (PN)-sensitized ALL and DS LAD2 cells under basal conditions and following PN (20 µg/mL) or anti-hIgE (10 µg/mL) stimulation (n = 3 independent experiments, each performed in quintuplicate). **(c)** Energy map under basal (left) and induced (right) conditions in ALL and DS cells following PN (20 µg/mL) or anti-hIgE (10 µg/mL) stimulation. The x- and y-axes represent ATP production derived from glycolysis (GlycoATP) and mitochondrial respiration (MitoATP), respectively (n = 2 independent experiments, each performed in quintuplicate). Data are presented as mean ± SEM.

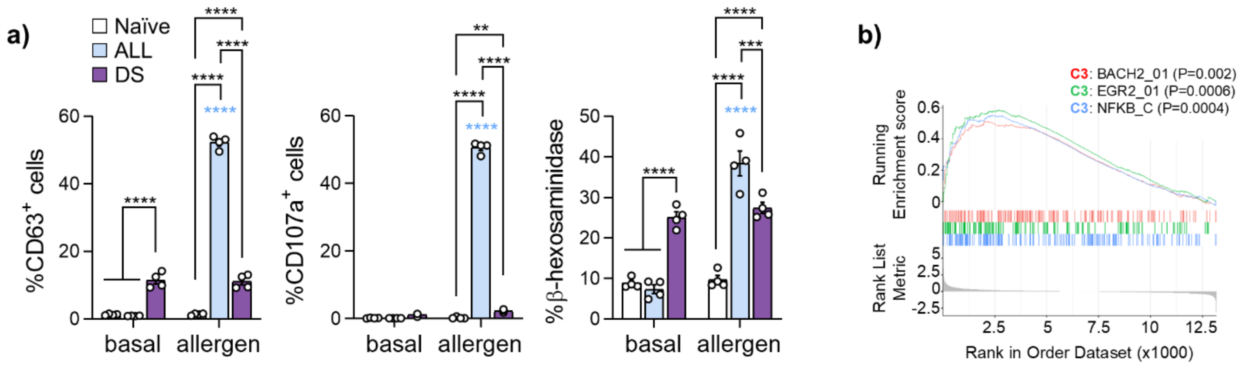


**Supplementary figure 5**. **Functional validation of transcriptional samples and regulatory gene enrichment. (a)** Degranulation analysis (CD63 and CD107a protein expression, and β-hexosaminidase release) of the four biological replicates of naïve, peanut (PN)-sensitized (ALL), and desensitized (DS) LAD2 cells used for RNA sequencing (assessed 2 h after desensitization). **(b)** Gene set enrichment analysis (GSEA) of DS cells using the regulatory target gene set (C3) collection of terms across all expressed genes (n = 13,168), ranked by $\log_2$ fold change *vs.* ALL cells. Data are presented as mean ± SEM. Statistical comparisons were performed using two-way ANOVA followed by Bonferroni multiple comparison test. Colored asterisks indicate significant differences relative to matched basal controls. **p < 0.01; ***p < 0.001; ****p < 0.0001.

### Supplementary tables

**Supplementary Table 1.** Clinical parameters of allergic sera used for mast cell experiments. Allergy diagnosis was confirmed by expert allergologists based on clinical criteria and evidence of sensitization.

| **Peanut allergic patients** | | | | | | |
|---|---|---|---|---|---|---|
| **Subject ID** | **Total IgE (kU/L)** | **Peanut (kU/L)** | **Ara h 1 (kU/L)** | **Ara h 2 (kU/L)** | **Ara h 3 (kU/L)** | **Dilution for sensitization** |
| 1# | 1,249 | ND | 52.9 | >100 | 26.6 | 1:200-1:100 |
| 2# | 179 | 54.9 | 20.4 | 24.5 | ND | 1:50 |
| 3* | 1,572 | ND | 34.1 | 44.5 | 46.9 | 1:50 |
| 4* | 974 | ND | >50 | >50 | 46 | 1:100 |
| 5* | 248 | ND | 12.55 | 8.5 | 7.7 | 1:50 |
| **Egg allergic patients** | | | | | | |
| **Subject ID** | **Total IgE (kU/L)** | **Egg white (kU/L)** | **Egg (kU/L)** | **Gal d 2 (kU/L)** | **Gal d 1 (kU/L)** | **Dilution for sensitization** |
| 6# | ND | ND | ND | 256 | ND | 1:25 |
| 7# | 287 | 55.1 | 81.9 | ND | 29.5 | 1:10 |
| 8* | >2,500 | 26.45 | ND | 25.44 | 0 | 1:50 |
| **Milk allergic patients** | | | | | | |
| **Subject ID** | **Total IgE (kU/L)** | **Cows' milk (kU/L)** | **α-lact (kU/L)** | **β-lact (kU/L)** | **Casein (kU/L)** | **Dilution for sensitization** |
| 9* | >2,500 | 32.3 | 10.11 | 7.51 | 6.76 | 1:25 |
| 10* | 1,512 | >50 | 49.7 | 25.7 | >50 | 1:50 |
| 11* | >2,500 | 3.8 | 12.6 | 32.6 | 5.8 | 1:25 |
| 12# | 2,165 | 635 | ND | ND | 658 | 1:50 |
| **Pollen allergic patients** | | | | | | |
| **Subject ID** | **Total IgE (kU/L)** | ***P. pratense* (kU/L)** | **Phl p 1 (kU/L)** | **Phl p 5b (kU/L)** | **Phl p 7 (kU/L)** | **Dilution for sensitization** |
| 13# | 1,554 | >100 | 47.3 | 52.2 | 0.03 | 1:50 |
| 14* | 1,062 | ND | 8.87 | 34.5 | ND | 1:100 |
| 15* | 137 | ND | 22 | 39.1 | ND | 1:100 |
| **Cat allergic patients** | | | | | | |
| **°Subject ID** | **Total IgE (kU/L)** | **Cat dander (kU/L)** | **Fel d 1 (kU/L)** | **Fel d 2 (kU/L)** | **Fel d 4 (kU/L)** | **Dilution for sensitization** |
| 13# | 1,554 | >100 | ND | 256 | ND | 1:50 |
| 16# | ND | ND | 21.4 | ND | 0.01 | 1:25 |
| 17# | 708 | 15.6 | 23.3 | 0.02 | 0.06 | 1:50 |

Abbreviations: lact = lactalbumin; ND = not determined; *P. pratense* = *Phleum pratense*. #ImmunoCAP (Phadia) determinations. Limits (kU/L): >17 = very positive; 3.5-17 = positive; 0.7-3.5 = moderate; 0.35-0.7 = mild; <0.35 = negative. *ALEX (Allergy Xplorer) determinations. Limits (kU/L): >15 = very high; 5-15 = high; 1-5 = moderate; 0.3-1 = low; <0.3 = negative.

**Supplementary Table 2**. List of antibodies used for flow cytometry and Western blot.

| REAGENT | CLONE | DILUTION | SOURCE | REFERENCE |
|---|---|---|---|---|
| **Flow Cytometry Antibodies** | | | | |
| **Activation panel** | | | | |
| Human specificity | | | | |
| Mouse anti-FcεRIα, FITC | AER-37 | 1:100 | BioLegend | #334608 |
| Mouse anti-CD117, PE | 104D2 | 1:600 | BioLegend | #313204 |
| Mouse anti-CD107a, APC | H4A3 | 1:100 | BioLegend | #328620 |
| Mouse anti-IgE, PE-Cy7 | MHE-18 | 1:100 | BioLegend | #325510 |
| Mouse anti-CD63, BV421 | H5C6 | 1:200 | BioLegend | #353030 |
| Mouse anti-CD63, PacB | H5C6 | 1:300 | BioLegend | #353012 |
| Murine specificity | | | | |
| Rat anti-IgE, PE | RME-1 | 1:200 | BioLegend | #406908 |
| Rat anti-CD107a, PerCP-Cy5.5 | 1D4B | 1:200 | BioLegend | #121625 |
| Rat anti-CD63, APC | NVG-2 | 1:200 | BioLegend | #143906 |
| Armenian hamster anti-FcεRIα, PE-Cy7 | MAR-1 | 1:200 | BioLegend | #134318 |
| Rat anti-CD117, BV421 | ACK2 | 1:200 | BioLegend | #135124 |
| Rat anti-CD117, PacB | 2B8 | 1:200 | BioLegend | #105820 |
| **Spleen co-culture panel** | | | | |
| Murine specificity | | | | |
| Rat anti-CD117, PacB | 2B8 | 1:200 | BioLegend | #105820 |
| Armenian hamster anti-FcεRIα, PE-Cy7 | MAR-1 | 1:200 | BioLegend | #134318 |
| Rat anti-CD44, BV785 | IM7 | 1:200 | BioLegend | #103041 |
| Rat anti-CD4, BV711 | GK1.5 | 1:200 | BioLegend | #100447 |
| **Western-blot Antibodies** | | | | |
| Rabbit monoclonal anti-phospho-Lyn (Y507) | 5B6 | 1:50,000 | Abcam | #ab278639 |
| Rabbit polyclonal anti-phospho-Src Family (Y416) | NA | 1:1,000 | Cell Signaling | #2101 |
| Rabbit monoclonal anti-phospho-LAT (Y200) | Y109 | 1:10,000 | Abcam | #ab68139 |
| Rabbit monoclonal anti-phospho-ERK1/2 (T202/Y204) | D13.14.4E | 1:2,000 | Cell Signaling | #4370 |
| Rabbit monoclonal anti-phospho-Akt (S473) | D9E | 1:2,000 | Cell Signaling | #4060 |
| Rabbit polyclonal anti-phospho-PLCγ (Y783) | NA | 1:1,000 | Cell Signaling | #2821 |
| Rabbit polyclonal anti-phospho-SHIP1 (Y1022) | NA | 1:1,000 | Cell Signaling | #3941 |
| Mouse monoclonal anti-β-actin | 8H10D11 | 1:1,000 | Cell Signaling | #3700 |

Abbreviations: NA = not applicable; FITC = fluorescein isothiocyanate; PE = phycoerythrin; PerCP-Cy5.5 = peridinin-chlorophyll-protein complex-cyanine 5.5; APC = allophycocyanin; PE-Cy7 = phycoerythrin-cyanine 7; BV = brilliant violet; PacB = pacific blue.

**Supplementary Table 3**. List of primers used for real time-qPCR.

| GENE (GeneBank accession) | PRIMER 5'-3' (Forward/Reverse) | AMPLICON SIZE | REFERENCE |
|---|---|---|---|
| **Target genes** | | | |
| In-home designed primers with Primer-BLAST | | | |
| ***CCL1*** (NM_002981.2) | GGCTCATCAAAGCTGCTCCA<br>AAGGGTACCTGCATGCTCTTG | 135 | |
| ***XIRP1*** (NM_194293.4) | GAGGAACGACGAACCCTGAG<br>GTAGAGGCTGGATGCAAGGG | 85 | |
| ***CHST1*** (NM_003654.6) | GTCACTTCGGTGTGGTTGGA<br>CAAGGGGTGAGGTCAAAGAGG | 188 | |
| ***GNG12*** (NM_018841.6) | CACCGGCAGGCGGATTC<br>GGTGCTTGCTGTTTTGCTGG | 113 | |
| ***RAI2*** (NM_001172732.2) | TCCACTGACCTGGTGGAGG<br>CAGGACCAGAGGGGAGTTGA | 147 | |
| ***TMEM151A*** (NM_153266.4) | CTGCGGGAAGAGCAGCG<br>GGATGAGCAGCGTGAGGAG | 94 | |
| ***PXDC1*** (NM_183373.4) | GGAGTTCTTCGAGATCCGCAC<br>TCCTTTATGGCAACCAGTCCTTG | 168 | |
| ***KCTD15*** (NM_001129994.2) | GGGGCTTCCAACGATACTCTG<br>GGACATGTTTCCTCCCTCCG | 124 | |
| ***SRRM3*** (NM_001110199.3) | GAAGAAGCATCGCCGAGACA<br>GGAGCTTCGAGATCTGTGCC | 70 | |
| ***RASD2*** (NM_001366725.1) | CGAAGGAGCAGCCCCGA<br>GAAGTCCTCGATGGTGGGTG | 185 | |
| ***RRAD*** (NM_001128850.2) | CGACTCAGACGAGAGCGTTT<br>CGATCATAGGTGTGCCCTGC | 132 | |
| ***EGR3*** (NM_001199880.2) | CGCGGTGGGAGAGAGAATG<br>GTTGGAAGGGGAGTCGAAGG | 147 | |
| ***MAFA*** (NM_201589.4) | AGAGCGAGAAGTGCCAACTC<br>GTACAGGTCCCGCTCTTTGG | 83 | |
| Primers obtained from literature | | | |
| ***IL27RA*** | TGCAGGTGAGCTACAAAGTCTGGT<br>AGCAGACCAAAGAGAGGTTGGTGA | 165 | PMID: 35117826 |
| ***CCL4L2*** | CCGCCTGCTGCTTTTCTTAC<br>TTGCTTGCCTACCACAGC | 107 | PMID: 34516352 |
| ***NOTUM*** | CTACTGGTGGAACGCAAACATGG<br>CGCACCACCTCCTGGATGATG | 129 | PMID: 34352447 |
| ***GZMB*** | GGTGGCTTCCTGATACAAGACG<br>GGTCGGCTCCTGTTCTTTGAT | 108 | PMID: 38607279 |
| ***PRRX2*** | AGCACAGTGCCACCCTACAG<br>CTTGGCAGGCTCTTCCACC | 383 | PMID: 11063257 |
| ***TEAD3*** | GCTCATTCCGCACATTCC<br>TATTGTGCTGGTTGCTCTGG | 172 | PMID: 24623739 |
| ***CXXC5*** | CAAGAAGAAGCGGAAACGCTGC<br>TCTCCAGAGCAGCGGAAGGCTT | 158 | PMID: 40001144 |
| **Housekeeping genes** | | | |
| Primers obtained from literature | | | |
| ***GAPDH*** | CGACCACTTTGTCAAGCTCA<br>CCCTGTTGCTGTAGCCAAAT1 | 59 | PMID: 17545502 |
| ***R18S*** | TCAACTTTCGATGATGGTAGTCGCC<br>TCCTTGGATGTGGTAGCCGTTT | ND | PMID: 35117826 |

Abbreviations: ND = not detected.

**Supplementary Table 4**. Summary of RNA-seq processing and pseudoalignment metrics.

| ID | CONDITION | N_PROCESSED READS | N_PSEUDOALIGNED READS | N_UNIQUE READS | P_PSEDOALIGNED READS | P_UNIQUE READS |
|---|---|---|---|---|---|---|
| A1 | Naïve | 24,018,296 | 21,299,828 | 5,833,056 | 88.7 | 24.3 |
| A2 | Naïve | 24,131,821 | 21,068,176 | 5,839,703 | 87.3 | 24.2 |
| A3 | Naïve | 21,972,425 | 19,261,383 | 5,308,321 | 87.7 | 24.2 |
| A4 | Naïve | 24,061,887 | 21,169,430 | 5,735,069 | 88.0 | 23.8 |
| B1 | ALL | 24,087,744 | 20,983,662 | 5,706,167 | 87.1 | 23.7 |
| B2 | ALL | 21,277,917 | 18,429,231 | 5,259,268 | 86.6 | 24.7 |
| B3 | ALL | 23,655,427 | 20,743,044 | 5,842,539 | 87.7 | 24.7 |
| B4 | ALL | 20,948,260 | 18,411,078 | 5,194,505 | 87.9 | 24.8 |
| C1 | DS | 21,539,781 | 18,732,766 | 5,242,008 | 87.0 | 24.3 |
| C2 | DS | 22,328,185 | 19,560,811 | 5,513,638 | 87.6 | 24.7 |
| C3 | DS | 21,663,155 | 18,912,535 | 5,416,852 | 87.3 | 25.0 |
| C4 | DS | 21,605,590 | 18,923,078 | 5,262,383 | 87.6 | 24.4 |

Abbreviations: N = number; P = percentage; ALL = allergic; DS = desensitized.
